# Transfer of Septin Rings to Cytokinetic Remnants Directs Age-Sensitive ER stress Surveillance Cell Cycle Re-entry

**DOI:** 10.1101/698829

**Authors:** Jesse T. Chao, Francisco Piña, Masayuki Onishi, Yifat Cohen, Maya Schuldiner, Maho Niwa

## Abstract

During cell division, cells must actively pass on organelles. Previously, we discovered the endoplasmic reticulum (ER) stress surveillance (ERSU) pathway that ensures the inheritance of functional ER. Activation of the ERSU causes the septin ring to mislocalize, which blocks ER inheritance and cytokinesis. Here, we found that the septin ring mislocalizes to previously utilized cell division sites called cytokinetic remnants (CRMs). The transfer of the septin ring to CRMs requires Nba1, a negative polarity component that normally prevents septin ring formation at CRMs. Furthermore, septin ring movement to CRMs relies on the ERSU component Slt2, which is recruited by binding Bem1. During ER stress, Bem1 also binds the GTP exchange factor Cdc24, without activating Cdc42, a GTPase that normally establishes polarized growth. Failure to translocate septin rings to CRMs delays the cell’s ability to re-enter cell division when ER homeostasis is re-established. Thus, ER stress considers the history of previous cell cycle for future cell cycle re-entry upon ER stress recovery.

## INTRODUCTION

The functions of eukaryotic cells are organized and distributed into specific organelles. During the cell cycle, not only does the genome divide but organelles must be correctly distributed. Thus, one of the fundamental questions in cellular biology is how specific organelles are inherited during the cell cycle. The endoplasmic reticulum (ER) is the major organelle responsible for the production and quality control of almost all secretory proteins. In addition, the ER is crucial for lipid biosynthesis and calcium homeostasis (Denic et al., 2006; Feige and Hendershot, 2011; Frakes and Dillin, 2017; Mori, 2000; Ron and Walter, 2007; Rutkowski and Kaufman, 2004). Importantly, the ER cannot be synthesized *de novo* and, instead, must be inherited from the mother cell. This suggests the presence of ER inheritance checkpoints.

The budding yeast, *Saccharomyces cerevisiae,* is an ideal model organism to study the division of the ER during the cell cycle, due to the asymmetric nature of yeast cell division. The ER in yeast is spatially segregated into the cortical ER (cER) that lies in the cell cortex, with some sections of the cER contacting the plasma membrane. The cER is connected by a few ER tubules with the peri-nuclear ER that is contiguous with the outer nuclear envelope (Barlowe, 2010; Bechmann et al., 2012; Du et al., 2004; Fehrenbacher et al., 2002; Hereford and Hartwell).

Previously, we discovered a cell cycle checkpoint for ensuring that functional ER is transferred to the daughter cell during the cell cycle, which we termed the ER stress surveillance (ERSU) pathway (Babour et al., 2010; Pina et al., 2018; Pina et al., 2016; Pina and Niwa, 2015). If the accumulation of unfolded or misfolded proteins exceeds ER functional capacity, ER homeostasis is disrupted, leading to a condition known as ER stress. In response to ER stress during the cell cycle, the ERSU pathway blocks the inheritance of the ‘*stressed ER’* into the daughter cell and mobilizes the septin ring from the bud neck, ultimately leading to cell cycle arrest at cytokinesis.

Surprisingly, the ERSU is independent of the well-known unfolded protein response (UPR) signaling pathway that regulates ER functional homeostasis. In parallel to the ERSU pathway, ER stress activates the UPR pathway to up-regulate the transcription of genes coding for ER chaperones and protein folding components to re-establish ER functional homeostasis (Ron and Walter, 2007). When ER functional homeostasis is re-established, cells are released from cell cycle arrest, re-enter the cell cycle, and inherit a functional ER into the daughter cell (Babour et al., 2010). Thus, the ERSU is one of the first cell checkpoint mechanisms, and works in concert with the UPR pathway to ensure proper organelle inheritance.

One of the hallmark events of the ERSU is septin ring mislocalization away from the bud neck. This process is a key mechanism leading to cytokinesis arrest in response to ER stress. The septin ring is composed of five septin subunits, Shs1, Cdc3, 10, 11, and 12, and its formation is dynamically regulated during the cell cycle (Field and Kellogg, 1999; Gladfelter et al., 2001; Mostowy and Cossart, 2012; Oh and Bi, 2011; Versele and Thorner, 2005; Weirich et al., 2008). Septin ring formation occurs even prior to the emergence of the daughter cell (bud), marking the incipient bud site. The formation of the septin ring is tightly linked to the targeting of activated Cdc42, which allows emergence and polarized growth of the bud. Cdc42 activity is tightly regulated by its upstream components including Cdc24, a GTP exchange factor (GEF) of Cdc42, and Bud1/Rsr1. The septin ring stays at the bud neck between the mother and daughter cells throughout most of the cell cycle. At the end of the cell cycle, the dividing membrane between the mother and daughter cell (i.e., the septum) forms, followed by cell division. Finally, the septin ring disassembles into subunits, which then re-initiate the cycle of assembly/disassembly. Interestingly, septin ring assembly is normally inhibited at cytokinesis remnants (CRMs), which are previously utilized cell cycle division sites, through a block involving Nba1 (Meitinger et al., 2014). This ensures that budding and subsequent cytokinesis occurs only at naïve locations that have never been used as cell division sites. The number of CRMs increases as a yeast cell undergoes more rounds of cell cycle, and thus it provides a molecular clock for the age of the yeast cell (Caudron and Barral, 2009).

Septin ring mislocalization is important for the ERSU pathway. In *slt2Δ* cells that are unable to mount the ERSU response, the septin ring remains present at the bud neck during ER stress and cells subsequently die. However, we still do not know the details of how septin rings become mislocalized, or the functional significance of this process. Therefore, in this study we used molecular and cell biological approaches to investigate septin ring dynamics during ER stress.

## RESULTS

### The septin ring moves to the bud scar in response to ER stress

To characterize septin ring movement during ER stress, we performed time-lapse microscopy. We monitored the morphology and localization of the septin ring in a wild-type (WT) cell carrying a green fluorescent protein (GFP)-tagged form of the Shs1 septin subunit (Shs1-GFP). Previously, we found that the targeting of septin subunits and the formation of septin rings occurred normally at the bud neck even under ER-stressed conditions (Babour et al., 2010). Thus, we reasoned that mislocalization of the septin ring must occur at a later stage of the cell cycle. We monitored the behavior of Shs1-GFP in large budded G2 cells. Under normal growth, as cells entered mitosis, the Shs1-GFP septin ring split into two and the fluorescent levels started to fade. Subsequently, Shs1-GFP began to accumulate at the incipient bud site (Figure 1A, Movie S1), consistent with previously reported septin ring dynamics (Oh and Bi, 2011; Versele and Thorner, 2005).

**Figure 1.**
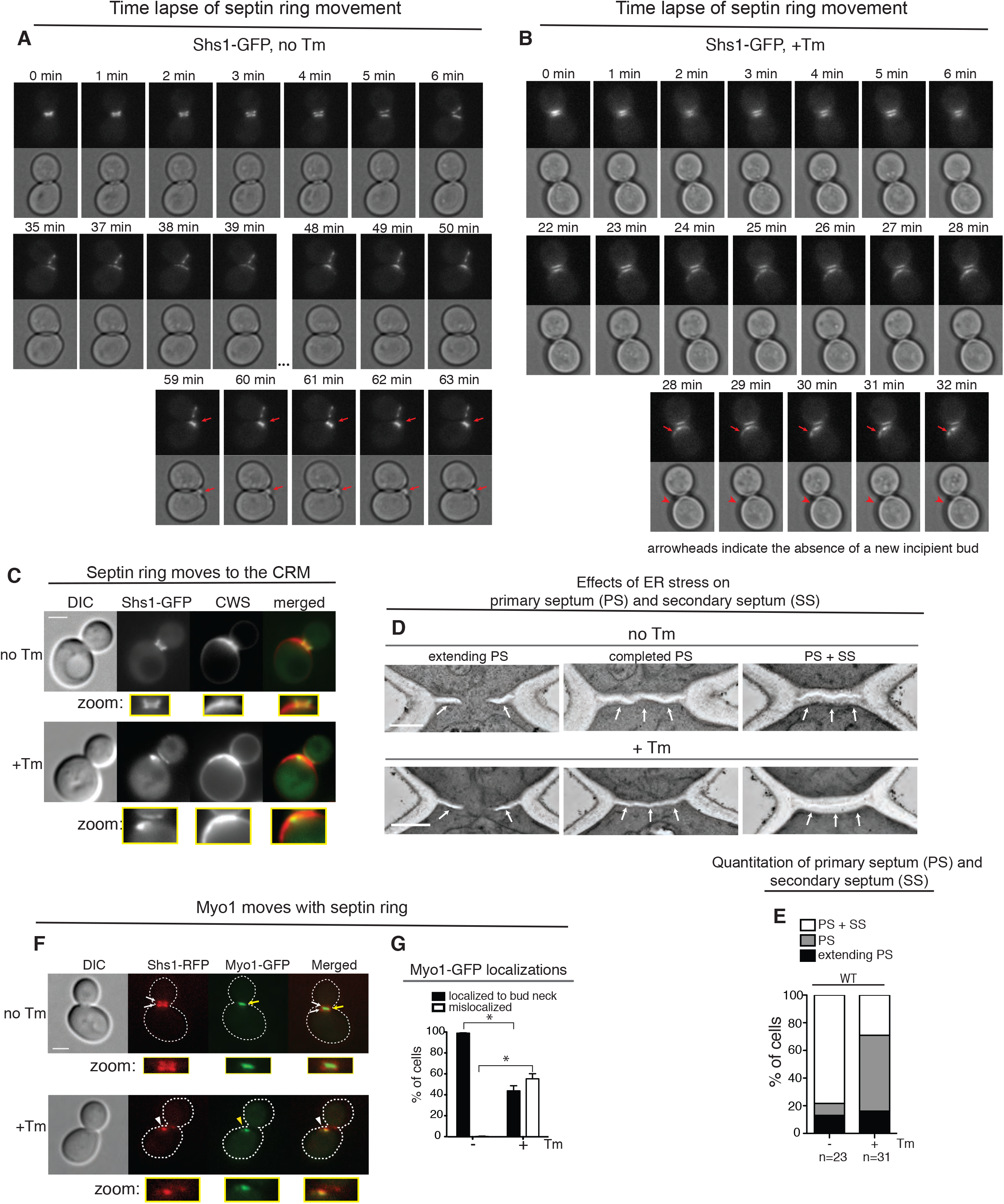
Septin rings relocalize to CRMs during ER stress. (A) Time-lapse analysis of septin dynamics (Shs1-GFP) in untreated WT cells. All scale bars, 2 µm. Later in the time course, a newly formed septin ring and a daughter cell are indicated by red arrows. See also Movie S1.
(B) Time-lapse analysis of septin rings in WT cells that were treated with 1 µg/ ml Tm. Red arrows point to translocated septin rings without the emergence of new buds, indicated by red arrowheads. See also Movie S2.
(C) Shs1-GFP cells were grown with (+) or without (-) 1µg/ ml Tm. CRMs were visualized by CWS. Inserts contain enlarged images cropped from the bud neck region.
(D) Representative electron micrographs taken from the bud neck regions of WT cells. Cells were synchronized for 30 min with alpha factor, released and either treated with DMSO (no Tm) or incubated with 15 µg/ml Tm for 2 hr (+Tm), and then fixed. White arrows indicate septums; scale bar represents 0.5 µm. PS, primary septum; SS, secondary septum.
(E) Quantification of the experiment in (D). Upon ER stress induction by Tm treatment, the number of cells with primary septum (PS) (but not secondary septum (SS)) (shown in gray) significantly increased. In contrast, both PS and SS formed in unstressed cells (white column).
(F) Myo1-GFP was co-localized with Shs1-RFP under both normal growth and ER stress. Cells were either grown without or with 1 µg/ml Tm. White arrows, RFP localization; yellow arrows, GFP. White and yellow arrowheads show altered localizations of Shs1 and Myo1, respectively.
(G) Quantification of Myo1-GFP in WT cells grown normally or under ER stress (1 µg/ml Tm) showed that Myo1-GFP moved with Shs1-RFP upon ER stress induction. Standard errors (SE) and statistic significances were calculated as described in materials and methods section unless otherwise stated.

To monitor Shs1-GFP dynamics under ER stress, we switched to growth medium containing tunicamycin (Tm), a well-characterized ER stress inducer. Tm inhibits protein N-glycosylation in the ER, resulting in an accumulation of unfolded proteins (Kuo and Lampen, 1974). In marked contrast to cells grown in normal media, the septin ring *transferred* to a site adjacent to the bud neck (Figure 1B, Movie S2). Importantly, we observed no apical growth of the new daughter cell at the site of septin translocation (Movie S2). We observed similar septin ring behavior when we activated ER stress by using *ero1-1* temperature-sensitive mutant cells at a non-permissive temperature (Figure S1A) (*ERO1* codes for the oxidoreductase that catalyzes disulfide bond formation in the ER. *ero1-1* at 37**°**C reliably induces the ER stress response (Frand and Kaiser, 1998; Pollard et al., 1998; Tu et al., 2000)). In addition, we found that the septin ring localization change in *ero1-1* at 37**°**C did not coordinate with the emergence of a new bud.

In haploid yeast of the BY genetic background, cells adopt an axial budding pattern in which new buds are consistently formed adjacent to the previous bud site or CRMs. This area includes both bud and birth scars, which can be visualized by calcofluor white staining (CWS). We found that Shs1-GFP co-localized with CRMs stained by CWS under ER stress (Figure 1C). We saw the same co-localization between another septin subunit, Cdc11-GFP, and CWS (Figure S1B). The septin translocation to CRMs was not specific to Tm, as we observed the same result in *ero1-1* cells at the non-permissive temperature (Figure S1A). Finally, we observed septin localization in CRMs in yeast cells of the W303 strain in which new buds emerge from distal positions with respect to the current site of division, presumably due to *BUD4* mutations (Figures S1C-S1E) (Voth et al., 2005). Therefore, we concluded that the septin ring moves to CRMs regardless of the type of ER stress or budding pattern.

### Septin transfer correlates with cytokinesis block during ER stress

As the septin ring is coordinated with cytokinesis and the formation of primary and secondary septum, which are cell walls that form in between mother and daughter cells (Onishi et al., 2013; Weiss, 2012; Wloka and Bi, 2012), we hypothesized that the translocation of the septin ring might disrupt septum formation. Electron microscopy analyses of both primary and secondary septum revealed that ER-stressed cells have significantly less completed secondary septum (Figures 1D and 1E). We also examined other components of cytokinesis such as Myo1, a type II myosin that forms an actomyosin contractile ring targeted to the bud neck by septins (Bi et al., 1998; Lippincott and Li, 1998; Tolliday et al., 2003). In unstressed WT cells carrying the genomic replacement of Myo1-GFP and Shs1-RFP, Myo1-GFP was localized at the bud neck sandwiched between two septin rings (Figure 1F). Upon ER stress induction, Myo1-GFP co-localized with transferred Shs1-RFP, and thus was no longer situated in the bud neck (Figures 1F and 1G). Taken together, these findings suggest that ER stress-induced septin movement leads to mislocalization of the actomyosin contractile ring, leading to a delay in cytokinesis. Curiously, *myo1Δ* cells can generate multi-budded and/or elongated cells due to the delayed cell division and separation despite continued polarized growth (Bi et al., 1998; Lord et al., 2005; Watts et al., 1987). The growth phenotype of ER-stressed cells differs from that of *myo1Δ* cells, suggesting that polarized growth is terminated in ER-stressed cells.

### Septin moves to bud scars and CRMs without polarized growth

Our finding that the transferred septin ring is recruited to CRMs during ER stress could partially explain the difference in phenotypes between *myo1Δ* cells and ER-stressed cells. Under normal growth, septin ring formation at the presumptive bud site is directed by the master regulator of cell polarity, Cdc42 (Okada et al., 2013). However, at CRMs, a Cdc42-inhibitory circuit is in place to prevent ‘refractory budding’ (Meitinger et al., 2014) from previously used bud sites. Thus, translocation of the septin ring to CRMs during ER stress could provide a mechanism to inactivate Cdc42 and block polarized growth.

In order to test if Cdc42 is inactivated during ER stress, we used a fluorescence biosensor, Gic2-PBD-RFP, to visualize only the active, GTP-bound form of Cdc42 (Okada et al., 2013). As Gic2 binds to only Cdc42-GTP, specific localization of Gic2 and its levels would reveal active Cdc42 levels. In unstressed cells, we found that active Cdc42 was enriched in the bud cortex, whereas the septin ring was localized in the bud neck (Figure 2A) as reported previously (Okada et al., 2013). In contrast, during ER stress, septin translocation coincided with the dispersal of active Cdc42 (Figure 2A), indicating that active Cdc42 did not accumulate at CRMs. Using time-lapse microscopy, we found that during normal growth, active Cdc42 accumulated in the growing bud of small-budded cells (Figure S2A). In contrast, in time-lapse imaging of a stressed cell with a smaller bud size, fluorescent signals of active Cdc42 dispersed within ∼30 minutes (Figure S2B).

**Figure 2.**
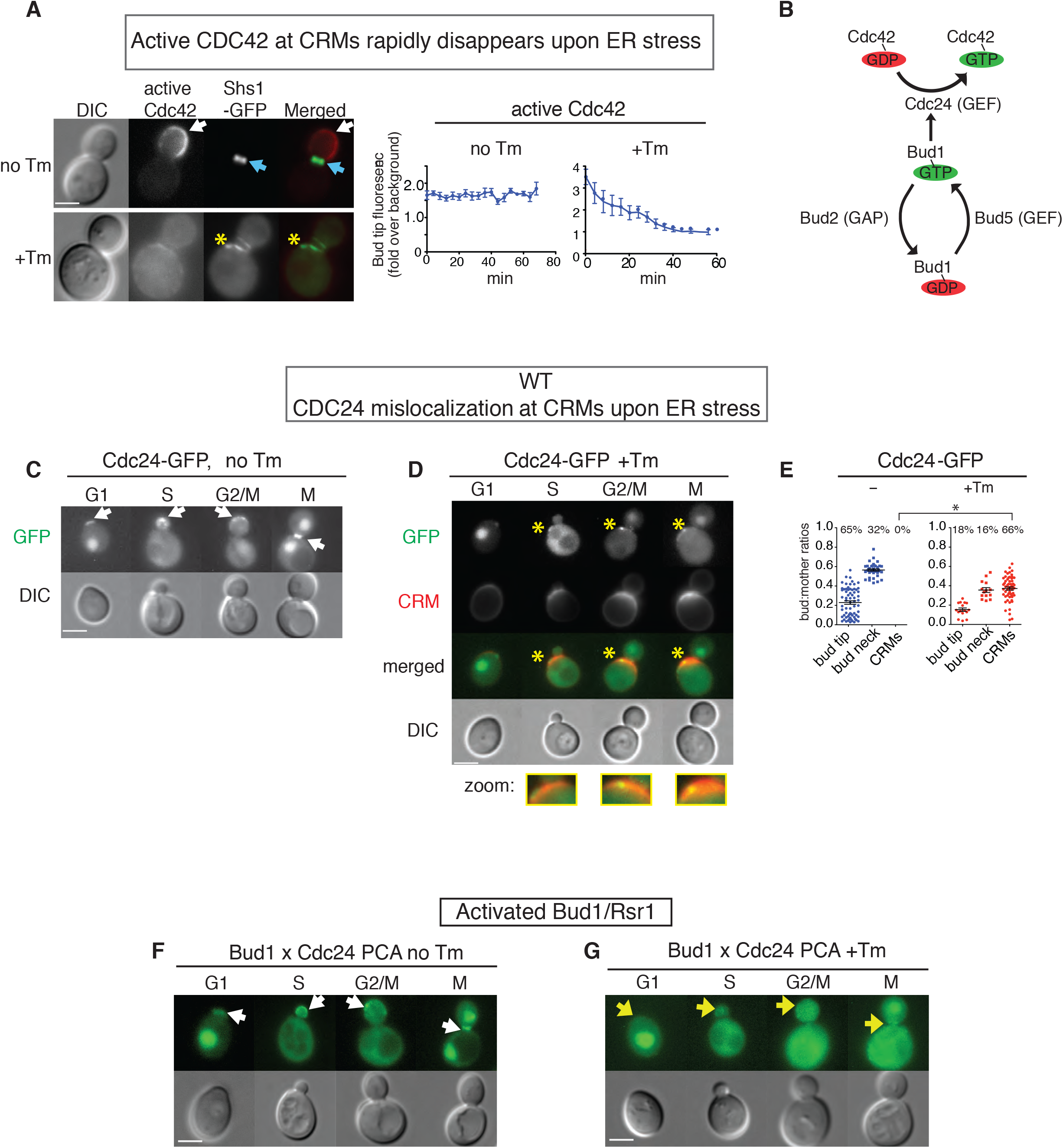
The Bud1/Rsr1 GTPase complex is negatively regulated during ER stress. (A) Cells co-expressing an active Cdc42 biosensor (Gic2-PBD-RFP) and Shs1-GFP were grown with or without 1µg/ml Tm. Left, locations of active Cdc42 are indicated by white arrows; Shs1-GFP is shown by blue arrows. Right, quantification of the average fluorescence intensities of active Cdc42 (by Gic2-PBD-RFP) at the bud tip in cells grown with or without 1 µg/ml Tm. Mislocalized Gic2-RFP is indicated with yellow asterisks. Measurements of Gic2-RFP levels were made for cells in each frame of time-lapse experiments shown in Figure S2B and S2C. See also Movies S3 and S4. All scale bars, 2µm.
(B) Schematics of the Cdc42-activating Bud1/Rsr1 GTPase module.
(C, D) Localizations of Cdc24-GFP in different stages of the WT cell cycle. (C) Cells that were untreated; (D) cells were treated with 1 µg/ml Tm. Cdc24 at the bud tip or bud neck is shown by white arrows; mislocalized Cdc24-GFP is indicated with yellow asterisks. CRMs were also visualized by staining with calcofluor white. In all microscopy panels, representative cells from each stage of the cell cycle are shown.
(E) Quantification of the Cdc24-GFP localizations shown in (C) and (D). The y-axis represents the ratios between the surface areas of buds and mothers; higher values indicate larger buds. The percentages of cells with the indicated localizations are shown above each column. Individual data points, as well as mean and SE are plotted, and the *p* value comparing localization of Cdc24-GFP to CRMs in untreated and treated samples was calculated. *, p < 0.0001.
(F, G) Split-YFP PCA between Bud1 and Cdc24. Representative cells from each stage of the cell cycle are shown. Cells were grown with or without 1 µg/ml Tm. White arrowheads in untreated cells (F) indicate sites where Bud1 interacted with Cdc24 and a yellow arrowhead in (G) shows a small amount of Bud1 interaction with Cdc24 in ER-stressed cells.

### ER stress disperses Cdc42 from the site of polarized growth, disconnecting its upstream effectors

To investigate the mechanism of Cdc42 inactivation from the site of polarized growth upon ER stress induction, we examined an upstream component, Cdc24, the GEF for Cdc42 (Figure 2B) (Hereford and Hartwell, 1974). In normal growing cells, Cdc24-GFP localized to sites of polarized growth including the incipient bud site in G1, the bud cortex in S and G2, and the bud neck in M phase (Figure 2C), as previously reported (Bos et al., 2007). We also detected Cdc24-GFP in the nucleus of G1 and M phase cells. In contrast, during ER stress the majority (66%) of cells had Cdc24-GFP translocated to CRMs (Figures 2D-2E), even though the nuclear localized Cdc24-GFP still remained during G1 and M phase.

Bud1/Rsr1 interacts with, and activates, Cdc24 at the polarized site of growth. Thus, we examined the active (GTP-bound) form of Bud1 using a split-yellow fluorescent protein (YFP) protein complementation assay (PCA) *in vivo*. In this assay, the YFP protein is split into two non-fluorescent fragments and each fragment is attached to a protein of interest. The interaction of these two proteins brings together two YFP fragments and restores YFP fluorescence. Under normal growth, we observed YFP signals generated from Bud1-Cdc24 interactions at the bud tip and bud neck (Figure 2F, white arrowheads), consistent with previous reports (Park et al., 1997; Park et al., 2002). During ER stress, however, we did not detect significant YFP signal either at the bud tip or the bud neck (Figure 2G: yellow arrowheads for loss of or reduced YFP signals). Together, our results suggest that while some Cdc24 remained at the bud neck, the majority no longer interacted with Bud1 (Figure 2F-2G).

We next examined the Bud1-activating components Bud5 (a GEF for Bud1) and Bud2 (a GTPase-activating protein, GAP, for Bud1) (Figure 2B) (Marston et al., 2001; Nelson et al., 2012). Specifically, we tested the Bud1/Rsr1-associated population of Bud5 (Figure S2D-S2E) and Bud2 (Figure S2F-S2G) using split YFP assays between Bud1-Bud5 and Bud1-Bud2, respectively. YFP generated from Bud1-Bud5 was localized at the bud tip (G1 and S) and the bud neck (G2/M and M) (Figure S2D). Similarly, YFP generated from Bud1-Bud2 was localized at the bud tip (G1 and S) and the bud neck (Figure S2F), consistent with previous report for localizations for Bud5 and Bud2 (Kang et al., 2001). In contrast, neither Bud1-Bud5 nor Bud1-Bud2 was localized at the site of polarized growth under ER stress (Figure S2E and S2G; yellow arrow pointing the loss of interaction). Furthermore, we did not observe significant accumulation of fluorescent signal at CRMs, indicating the lack of Bud1-Bud5 and Bud1-Bud2 localization at CRMs. Taken together, these results show that ER stress mislocalizes Cdc24 to CRMs and attenuates the activity of the Bud1 GTPase.

### ER stress-induced removal of the septin ring from the bud neck, but not translocation to CRMs, is sufficient to inactivate Cdc42

We next examined whether septin ring translocation to CRMs is critical for inactivation of either Cdc42 or its activating components. To this end, we generated a septin mutant that can be mislocalized from the bud neck but does not reached to CRMs during ER stress: a C-terminal truncation of the Shs1 septin subunit, Shs1ΔCTE (CTE: C-terminal extension; Figure 3A). During normal growth, Shs1ΔCTE-GFP was localized at the bud neck like WT Shs1-GFP (Figure 3B). However, upon ER stress induction ∼70% of *shs1ΔCTE* cells had no detectable septin ring at CRMs (Figure 3B). Shs1ΔCTE-GFP expression levels were similar to WT Shs1-GFP even after ER stress induction (Figure 3C), revealing that septin rings were dispersed throughout the cytosol and that the Shs1 C-terminal fragment was responsible for septin ring formation at CRMs under ER stress.

**Figure 3.**
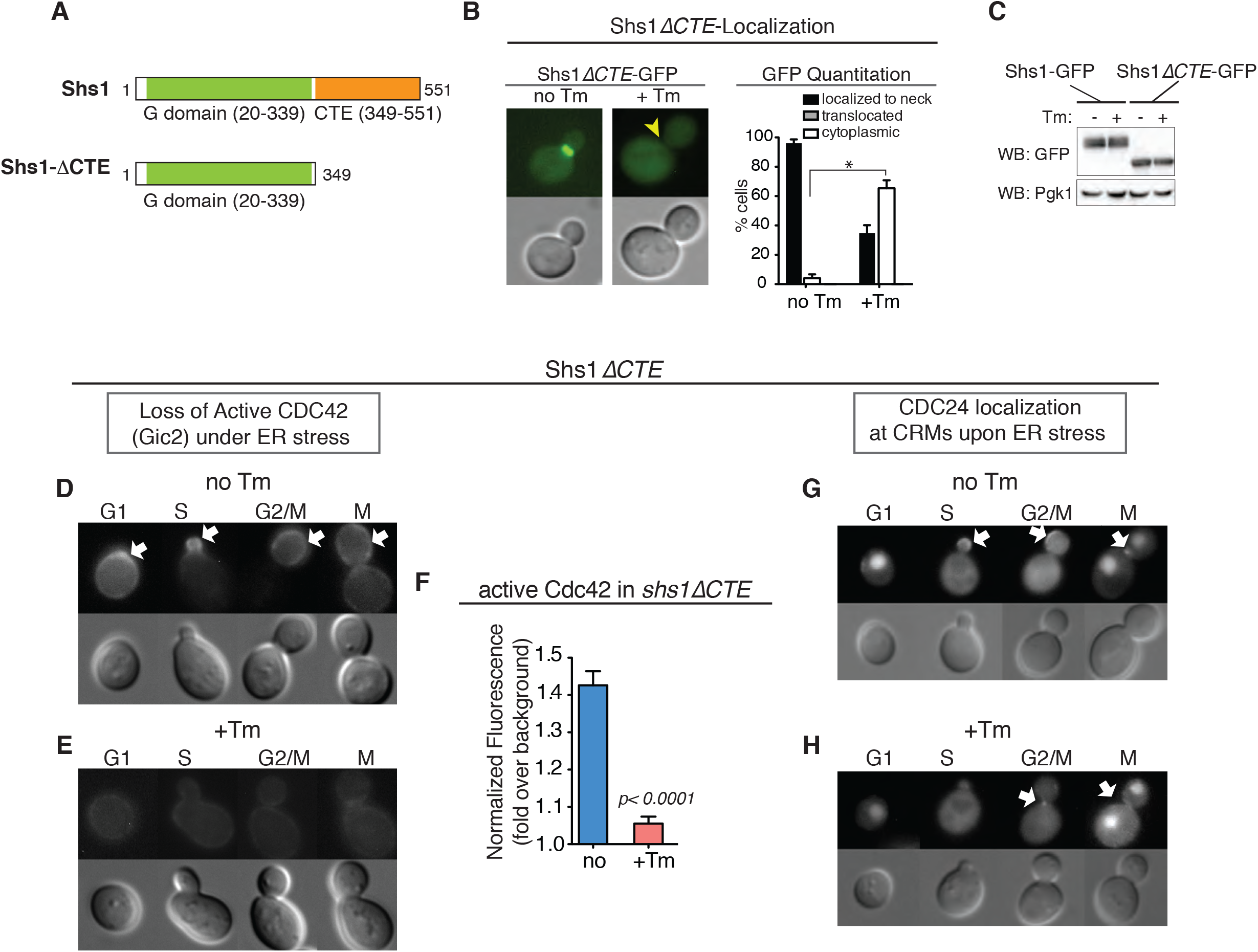
The CTE of Shs1 is required for septin ring translocation to CRMs in response to ER stress. (A) Domain organization of Shs1 (Versele and Thorner, 2004), the G domain, and GTPase binding. Shs1*ΔCTE* lacks C-terminal end (CTE) of WT Shs1 (aa. 349-551).
(B) Shs1*ΔCTE*-GFP is dispersed in cells grown with 1 µg/ml Tm, while it is localized at the bud neck during normal growth. Arrowheads point indicate absence of GFP localization. All scale bars, 2 µm. Quantification of Shs1*ΔCTE*-GFP in WT cells with or without 1 µg/ml Tm. * represents *p*<0.0001 comparing % of cells with dispersed Shs1*ΔCTE*-GFP.
(C) Western blot analysis of Shs1-GFP and Shs1Δ*CTE*-GFP expression in cells with or without 1 µg/ml Tm. Anti-Pgk1 was used as a loading control.
(D and E) Localization of Cdc42 (Gic2) in *shs1ΔCTE* cells that were either (D) untreated (no Tm) or (E) treated with 1µg/ml Tm. White arrows show the polarized localization of Gic2-GFP in unstressed cells (D), which was lost in ER-stressed cells.
(F) Quantification of active Cdc42-GFP fluorescence levels in *shs1*Δ*CTE* cells grown with or without 1 µg/ml Tm. Mean and SE were calculated based on values from at least three independent experiments.
(G and H) Localization of Cdc24-GFP in *shs1ΔCTE* cells that were either (G) untreated or (H) treated with 1µg/ml Tm. ER stress caused mislocalization of Cdc24-GFP.

In ER-stressed *shs1ΔCTE* cells, Cdc42 was no longer present at the bud tip or the bud neck, but rather was dispersed upon ER stress induction to a similar extent as ER-stressed cells with the WT Shs1 subunit (Figures 3D-3F). During ER stress, Cdc24 in *shs1ΔCTE* cells was also mislocalized similarly to WT cells (Figures 3G and 3H). Thus, despite the lack of septin ring assembly at CRMs in ER-stressed *shs1ΔCTE* cells, Cdc42 inactivation took place normally. This suggests that loss of the septin ring from the bud neck, but not formation of septin rings at CRMs, is sufficient to inactivate Cdc42 during ER stress.

### *shs1ΔCTE* cells are able to induce the ERSU pathway

We next explored whether translocation of the septin ring to the CRM is critical to block cER inheritance in response to stress. We assessed ER inheritance block by quantifying the number of cells without the cER in the daughter cell upon ER stress induction by Tm (Figure S3A). We classified cells as before to three groups: cells with small buds (< 2 μm; Group 1), cells with medium-sized buds but without nuclei (Group II), and large budded cells with nuclei (Group III) (Babour et al., 2010; Pina et al., 2016; Pina and Niwa, 2015). Previously, we demonstrated that ER stress does not block ER inheritance and cytokinesis in the first round of the cell cycle in cells that have already inherited the cER (some Class II and Class III cells), but it does block ER inheritance in the second round of the cell cycle (Pina and Niwa, 2015). Thus, with an asynchronous population of cells, the level of ER inheritance for class I cells represents most closely the impact of ER stress. Therefore, throughout this study, we focused on percent ER inheritance block of Class I cells as a representation of ER inheritance block. We found that ER stress blocked cER inheritance in WT Class 1 cells (Figure S3A, lanes 1-2) but not in ERSU-deficient *slt2Δ* cells (Figure S3A, lanes 13-14), as we reported previously (Babour et al., 2010; Pina et al., 2018). ER stress blocked inheritance of the cER in *shs1ΔCTE* cells at a level similar to that of WT cells (Figure S3A; lanes 1-2 for WT vs. lanes 7-8 for *shs1ΔCTE* cells). These results suggest that septin ring movement away from the bud neck is important for cell survival during ER stress, but septin ring movement to CRMs itself is not required. Thus, the functional significance of septin ring translocation to CRMs may reside beyond the initial stages of the ERSU pathway.

### Slt2 is required for ER stress-induced Cdc42 inactivation

The ability of *shs1ΔCTE* cells to inactivate Cdc42 in response to ER stress suggests that septin ring removal from the bud neck, but not transfer to CRMs, is required for inactivation of Cdc42. Furthermore, even though Cdc42 was inactivated, some levels of Cdc24 still moved to CRMs even in the absence of septin ring transfer at CRMs in *shs1ΔCTE* cells. These findings prompted us to address two important questions: what is the mechanism of Cdc42 inactivation? And, what is the functional significance of septin ring translocation to CRMs in response to ER stress?

We reported previously that Slt2 is a component of the ERSU pathway and is important for septin ring translocation (Figure S3B) and cER inheritance block (Figure S3A) (Babour et al., 2010). Thus, we tested if Slt2 is also required for Cdc42 inactivation during ER stress. In ER-stressed *slt2Δ* cells, Cdc42 remained at the site of polarized growth (Figures S3C and S3D). Furthermore, Cdc24 was unchanged in unstressed *slt2Δ* cells (Figures S3E-G), and ER stress induction did not result in its translocation (Figure S3E). Therefore, Slt2 plays a role in the inactivation of Cdc42 under ER stress in the ERSU pathway.

### Cdc42 inactivation status of *slt2Δshs1ΔCTE* cells

No previous study in *S. cerevisiae* has reported the involvement of MAP kinases in the regulation of Cdc42 during normal growth conditions. Thus, to further dissect the mechanism of Slt2-induced Cdc42 inactivation we tested if the loss of Slt2 in *shs1ΔCTE* cells also blocked the inactivation of Cdc42. In *slt2Δshs1ΔCTE* double mutant cells, we found that approximately 25% of *slt2Δ shs1ΔCTE* cells had two or more buds regardless of ER stress (Figures 4A), revealing that these cells had lost the ability to select a single polarity site (Chiou et al., 2017). We did not observe any multi-budded cells in either *slt2Δ* or *shs1ΔCTE* single mutants. Thus, the combined effects of deleting Slt2 and Shs1-CTE must contribute to this phenotype. This is consistent with the ability of cell polarity to affect cell division by impacting the cytokinesis machinery (Wu et al., 2013). Surprisingly, the septin ring (visualized by Cdc11-GFP) in multi-budded *slt2Δshs1ΔCTE* cells was localized at the bud neck, but became fragmented in response to ER stress (Figures 4B and 4C; single-budded cell).

**Figure 4.**
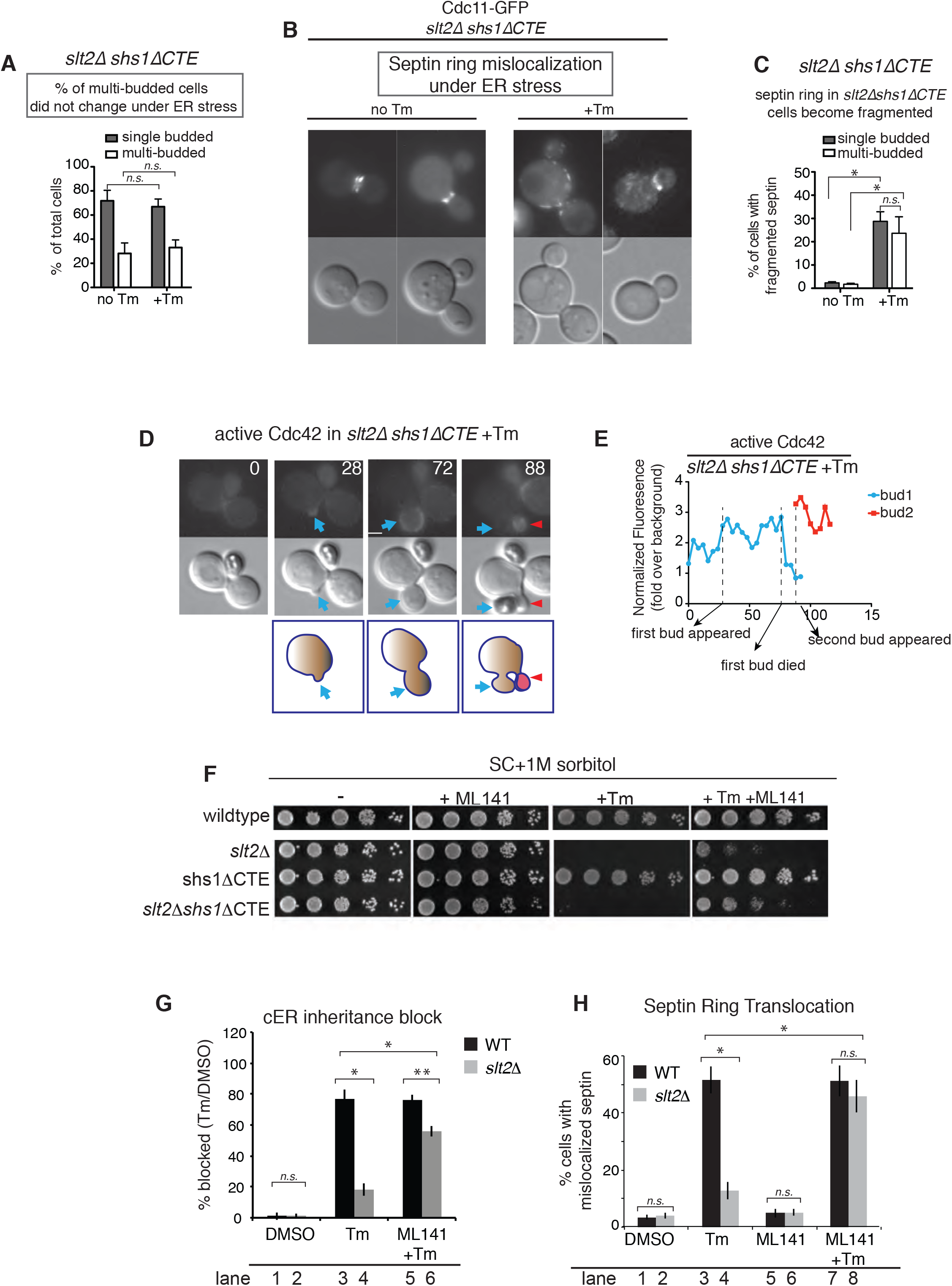
Cdc42 is inactivated by Slt2 in *Shs1ΔCTE* cells. (A, B) Some *slt2Δ shs1ΔCTE* cells were multi-budded even under normal growth without Tm. To visualize the septin ring, GFP-tagged Cdc11 (Cdc11-GFP), another septin subunit, was used (B). Cdc11-GFP was localized at the bud neck of single-budded *slt2Δ shs1ΔCTE* cells under normal growth conditions, but remained at the bud neck and did not translocate to CRMs upon ER stress induction (+Tm). However, some levels of mislocalization were detected in single-budded ER-stressed cells (A). In multi-budded *slt2Δ shs1ΔCTE* cells, Cdc11-GFP was localized only at one bud neck without ER stress, and it was mislocalized in the presence of Tm (B). % of single and double budded cells between no Tm and +Tm were not significantly different (n.s).
(C) Quantitation showed that septin ring fragmentation was increased in ER stressed both single-budded and multi-budded *slt2Δ shs1ΔCTE* cells. *represents *p* < 0.001 and *ns.,* not significant.
(D) Time-lapse analysis of active Cdc42 (Gic2-GFP) in the first and second buds emerged upon addition of Tm to *slt2Δ shs1ΔCTE* cells (time 0). Images from the time-lapse analysis taken at 0, 28, 72, and 88 sec are shown. Blue arrows show the first bud and red arrowheads show the second bud emerged from the original mother *slt2Δ shs1ΔCTE* cells.
(E) Quantification of Gic2-GFP fluorescence levels in the first (blue) bud and the second (red) bud at indicated times.
(F) Inactivation of Cdc42 rescued ER-stressed *slt2Δ* and *slt2Δ shs1ΔCTE* cells. A tenfold dilution of WT, *slt2Δ, shs1ΔCTE,* and *slt2Δ shs1ΔCTE* cells were spotted on synthetic complete medium with no Tm, 20µM ML141, +0.5 μg/ml Tm or +(0.5 μg/ml Tm+20µM ML141). ML141 is a well-characterized Cdc42 inactivating agent (ref). As Slt2 is known to be involved in the cell wall integrity response (Verna et al., 1997), 1M sorbitol was added to suppress the cell wall integrity response.
(G) cER inheritance block of the ER-stressed *slt2Δ* cells (lane 4) was restored upon treatment with ML141 (lane 6). In contrast, the inheritance of cER failed to occur in ER-stressed *slt2Δ* cells (lane 4). Data are the mean ±SD of three independent experiments; n>100 cells of each strain. Unpaired two-tailed t-tests values comparing WT and *slt2Δ* cells are shown: ns, not significant; *, *p* < 0.01; **, *p* < 0.05.
(H) Septin ring localization in either WT or *slt2Δ* cells, treated with DMSO, Tm (0.5 μg/ml), ML141 (20µM), or ML141 (20µM) plus Tm (0.5 μg/ml). The septin ring failed to translocate in ER stress-induced ERSU-deficient *slt2Δ* cells (lane 4), and ML141 treatment of ER-stressed *slt2Δ* cells restored septin ring translocation to CRMs (lane 8). Data are the mean ± SD of three independent experiments; n>100 cells of each strain. P-values from unpaired two-tailed t-tests comparing WT and *slt2Δ* cells are shown: ns, not significant; *, *p* < 0.01; **, *p* < 0.05.

We next examined Cdc42 in *slt2Δshs1ΔCTE* cells (Figures 4D and 4E, Movie S5). In time-lapse experiments of *slt2Δ shs1ΔCTE* cells, activated Cdc42 was localized at the bud tip (Figure 4D; blue arrowheads). However, following ER stress, a second bud started to emerge without cytokinesis of the first daughter cell, and Cdc42 moved to the cortex of the second daughter cell (Figure 4D; red arrowheads and Figure 4E; bud 1 (blue) to bud 2 (red)). Furthermore, the Cdc42 upstream effector Cdc24 moved to the second bud emergence site regardless of ER stress (Figures *S4A* and *S4B*). Thus, the loss of Slt2 in *shs1ΔCTE* cells disrupted Cdc42 inactivation in response to ER stress. Consistent with the multi-budded phenotype and the lack of Cdc42 inactivation, growth of *slt2Δ shs1ΔCTE* cells was significantly diminished under normal growth, and ER stress caused cell death (Figure 4F; +Tm).

### Inactivating Cdc42 can rescue ERSU-deficient *slt2Δ* cells under ER stress

The above results revealed an unprecedented and essential role of Slt2 MAP kinase in the inactivation of Cdc42 during ER stress. To further evaluate the functional significance of the Slt2-dependent inactivation of Cdc42, we tested whether inactivating Cdc42 via its inhibitor ML141 could rescue the growth of *slt2Δ* cells. Indeed, ML141 rescued the growth of both *slt2Δ* and *slt2Δ shs1ΔCTE* cells under ER stress (Figure 4F; compare +Tm vs. +Tm+ML141). Significantly, ML141 treatment also rescued the ER inheritance block (Figure 4G; compare lanes 4 and 6, Figure *S4D*) and septin transfer to CRMs (Figure 4H; compare lanes 4 and 8, Figure *S4C*) in ER-stressed *slt2Δ* cells. These results are consistent with the idea that Slt2-induced Cdc42 inactivation is a hallmark of the ERSU pathway, coordinating with other ERSU events such as ER inheritance block and septin ring movement to CRMs to ultimately contribute to cell survival in response to ER stress.

### Split-DHFR screen identifies Slt2 functional partners in the ERSU pathway

The importance of Slt2 is underscored by its involvement in a wide range of cellular functions, including genome silencing, cell wall responses, and polarized growth (Chen and Thorner, 2007; Gustin et al., 1998). As a result, Slt2 is localized throughout the cell. This makes it challenging to identify Slt2 that is specifically functioning in the ERSU pathway. To examine how Slt2 is involved in Cdc42 inactivation in response to ER stress, we performed a split-DHFR screen to quantitatively identify Slt2 binding partners *in vivo* (Tarassov et al., 2008). Split-DHFR is a growth-based selection assay, in which bait and prey proteins are tagged with complimentary fragments of a mutated DHFR (mDHFR); if bait and prey proteins interact, the two fragments of mDHFR reconstitute into a fully functioning enzyme that is not inhibited by the inhibitor of the endogenous and essential yeast DHFR (Tarassov et al., 2008). Therefore, growth levels on methotrexate are proportional to the amount of reconstituted mDHFR enzyme and enable quantification of protein-protein interactions. Using this assay, we screened ∼6,000 genes (Table S1) and identified 100 that showed enhanced interactions with Slt2 specifically during ER stress (Table S2).

### Bem1 is a novel binding partner for Slt2 during ER stress

We further conducted gene ontology (GO) analysis on the 100 identified genes and categorized them according to their GO molecular functions (Figure *S5A,* Table S3) (Robinson et al., 2002). We noticed that Bem1, Bnr1, and Vrp1 were the most connected genes (Figure *S5A*). Bem1 is an important polarity factor that functions as a scaffolding protein for Cdc24 and Cdc42, and recruits them to sites of polarized growth at both the bud neck and bud tip (Pruyne and Bretscher, 2000; Pruyne et al., 2004). Additionally, Bem1 is sequestered in CRMs by Nba1 and prevents it from binding to Cdc24, as a way of inhibiting Cdc42 recruitment to CRMs and of reducing the functional pool of Bem1 in cells (Meitinger et al., 2014). Thus, we focused on Bem1. Depending on which Bem1 pool Slt2 interacts with, we may be able to predict the functional significance of the Slt2-Bem1 interaction.

Using GFP-tagged Bem1, we found that Bem1 was localized at the bud tip in the early phase of the cell cycle and at the bud neck and CRMs later in the cell cycle under normal growth (Figure 5A), in agreement with previous reports (Liu and Novick, 2014; Madden and Snyder, 1998; Smith et al., 2013; Toenjes et al., 2004). Upon ER stress induction, we observed Bem1-GFP primarily at the bud neck and CRMs, and only a small amount of Bem1-GFP at the site of polarized growth (i.e., the bud tip) (Figure 5B). The loss of Bem1 localization at the bud tip is consistent with loss of activated Cdc42 (Gic2) from the bud tip and loss of polarized growth during ER stress (Figure 2A).

**Figure 5.**
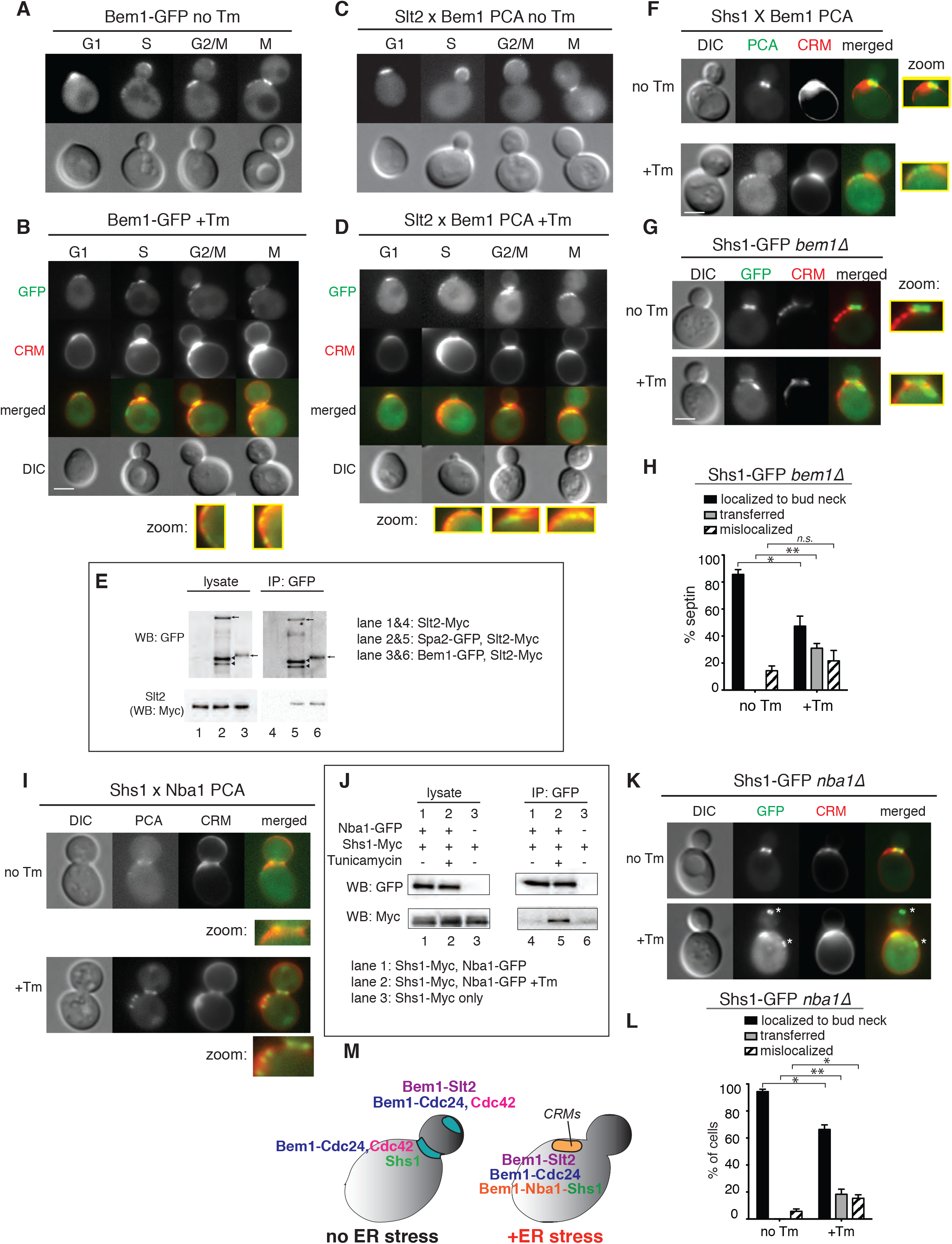
Bem1 interaction with Slt2 facilitates translocation of septin rings at CRMs during ER stress. (A) Bem1-GFP in unstressed WT cells was localized to the bud tip, bud neck, and CRMs.
(B) Bem1-GFP localization was altered in Tm-treated WT cells. CRMs were visualized by staining with WGA-594. Zoomed-in views of Bem1-GFP at CRMs are shown.
(C) Split-YFP PCA between Slt2 and Bem1 in untreated WT cells (no Tm).
(D) Split-YFP PCA between Slt2 and Bem1 in cells treated with 1µg/ml Tm. Close-up views of Slt2 interacting with Bem1 at CRMs are also shown.
(E) We performed co-IP experiments with Bem1-GFP and Slt2-Myc (lane 6). The extent of immunoprecipitation was similar to that of a previously identified Slt2 binding protein, Spa2-GFP (lane 5).
(F) Split-YFP PCA between Shs1 and Bem1 in untreated (no Tm) and Tm-treated WT cells. Close-up views show Shs1 interacting with Bem1 localized at the bud neck in untreated cells and at CRMs in Tm-treated cells. Zoomed-in views showing PCA signals at the bud neck for unstressed and at CRMs for ER-stressed cells are shown.
(G) Shs1-GFP in unstressed cells was localized to the bud neck of *bem1Δ* cells, whereas Shs1-GFP was mislocalized away from the bud neck but was localized outside of CRMs in Tm-treated *bem1Δ* cells. Zoomed-in views show Shs1-GFP at the bud neck in unstressed cells and at a location distinct from CRMs in ER-stressed cells.
(H) Quantification of Shs1-GFP in *bem1Δ* cells shows that significant levels of Shs1-GFP become mislocalized upon ER stress induction. *, *p* < 0.01; **, *p* < 0.001; *ns*, not significant.
(I) Split-YFP PCA between Shs1 and Nba1 in untreated (no Tm) and Tm-treated WT cells. Zoomed-in views show PCA signals at the bud neck in unstressed cells (no Tm) and at CRMs in ER-stressed cells (+Tm).
(J) We detected co-IP of Shs1-Myc and Nba1-GFP only after Tm treatment of WT cells (lane 5). In untreated cells, no physical interaction between Shs1-Myc and Nba1 was detected (lane 4).
(K) Nba1 is required for Shs1-GFP localization at CRMs during ER stress. Shs1-GFP in untreated (no Tm) or Tm-treated (+Tm) *nba1Δ* cells is shown.
(L) Quantification of Shs1-GFP in unstressed and ER-stressed *nba1Δ* cells. *, *p* < 0.05; **, *p* < 0.005.
(M) A schematic summarizing our findings on translocation of septin rings at CRMs during ER stress. Bem1-Cdc24 and Slt2 were localized at sites of polarized growth in unstressed cells; ER stress caused Bem1-Cdc24, Slt2, and Shs1 to translocate to CRMs. Shs1 localization at CRMs depended on Nba1 and Bem1.

These findings suggest that Slt2 localization to CRMs occurs by its association with CRM-localized Bem1. In order to visualize which pool of Bem1 interacts with Slt2, we employed split-YFP PCA in living cells (Michnick et al., 2007). During normal growth, Slt2 interacted with Bem1 at sites of polarized growth, namely at the bud tip in small budded cells indicative of S phase, and the bud neck in large budded cells indicative of G2/M phases (Figure 5C). We further verified the interaction between Slt2 and Bem1 biochemically by co-immunoprecipitation (co-IP). The amount of Slt2 co-purified with Bem1 was similar as Slt2 with Spa2, a protein known to interact with Slt2 (Figure 5E, compare lanes 5 and 6) (van Drogen and Peter, 2002). During ER stress, Slt2-Bem1 PCA was localized at CRMs (Figure 5D), suggesting that Slt2 interacts with the pool of Bem1 at CRMs.

### Bem1 recruits the septin ring to CRMs during ER stress

In addition to Slt2, the septin ring also mobilized to CRMs from the bud neck (Figures 1A and 1B). Thus, we tested if the septin ring interacts with Bem1 at CRMs during ER stress using a PCA assay. In unstressed cells, we observed fluorescent signals at the bud neck, revealing that Shs1 interacted with Bem1 at this location; however, during ER stress fluorescence signals representing the Shs1-Bem1 interaction were located at CRMs (Figure 5F). Shs1-GFP failed to accumulate on CRMs in ER-stressed *bem1Δ* cells (Figure 5G), revealing the importance of Bem1 for septin ring transfer to CRMs. In *bem1Δ* cells, Shs1-GFP was mislocalized at a location on the cell cortex outside of the CRM (Figures 5G and 5H).

### A negative polarity establishment factor, Nba1, remains at CRMs and is required for Shs1 localization to CRMs under ER stress

Our finding that Bem1 at CRMs binds to Slt2 and Shs1 was rather unexpected because Bem1 binding to Cdc24 at CRMs is normally prevented by Nba1, a recently identified CRM landmark (Meitinger et al., 2014). Under normal growth, Nba1 prevents both polarized growth at CRMs by blocking Bem1 binding to Cdc24. A genome-wide study revealed genetic interactions between Nba1 and Shs1 as well as components of the ERSU pathway, including Slt2, Bck1, and Pkc1 (Figure *S5B*) (Costanzo et al., 2011). This suggests that Nba1 may be an integral part of the ERSU pathway, and provides further rationale for testing Nba1.

Given our finding that Slt2 (Figure 5D), Shs1 (Figures 5F and 5G), and Cdc24 localize to CRMs under ER stress (Figure 2D), we next tested if Nba1 remained localized at CRMs in response to ER stress. We found that Nba1-GFP was localized to the bud neck and CRMs (Figure *S5C*), similar to previous reports (Meitinger et al., 2014). The localization of Nba1-GFP to CRMs did not change during ER stress (Figure *S5D*), consistent with Nba1’s role as a landmark for CRMs. Using the split-YFP assay, we found that Shs1-RFP and Nba1-GFP co-localized at CRMs only under ER stress (Figure 5I; +Tm). We also detected the Shs1-Nba1 interaction using co-IP, but the interaction was only significant in the presence of Tm (Figure 5J, compare lanes 5 (+Tm) vs 6 (-Tm)). Importantly, in *nba1Δ* cells, only 23% of cells showed a transferred septin ring in CRMs (Figures 5K and 5L), revealing a role for Nba1 in recruiting Shs1 to CRMs during ER stress (Figure 5M).

A recent study reported that two transmembrane proteins, Rax1 and Rax2, help to localize Nba1 at CRMs (Figure S5E) (Meitinger et al., 2014). We confirmed that Nba1 localization at CRMs indeed depended on Rax1, but its localization to the bud neck did not (Figures S5F and S5G). If bud neck-localized Nba1 is sufficient to target the translocated septin ring to CRMs during ER stress, we would not expect *RAX1* deletion to affect septin translocation. To differentiate which populations of Nba1 are responsible for mediating septin ring translocation during ER stress, we tested Shs1-GFP localization in ER-stressed *rax1Δ* cells. We found that Shs1-GFP remained at the bud neck, or mislocalized outside of CRMs in response to ER stress in *rax1Δ* cells (Figures S5H and S5I), indicating that Nba1 in CRMs is critical for septin ring translocation.

### The functional significance of Slt2 and Shs1 localization at CRMs

The above results revealed that components that decollate CRMs undergo significant changes under ER stress (Figure 5M): (1) Slt2 is localized at CRMs upon interaction with Bem1 (Figure 5D), (2) septin ring transfer to CRMs requires Bem1 (Figures 5F and 5G) and (3) Nba1 is required for septin transfer to CRM (Figures 5K and 5L). We next investigated the functional significance of septin ring transfer to CRMs by comparing WT and *shs1ΔCTE* cells. As ER inheritance block occurred normally in both WT and *shs1ΔCTE* cells (Figure S3A), we examined a later event: the ability of cells to re-enter the cell cycle after ER functional homeostasis is re-established. In order to test whether *shs1ΔCTE* cells are able to re-enter the cell cycle in a manner similar to WT cells after recovery from ER stress, we devised an ER stress-recovery assay (Figure 6A). In this method, we first treated the cells with Tm for 2 hrs, and then washed them quickly in warm YPD medium without Tm before starting the recovery time course. At 4 min after Tm washout and recovery from ER stress, the septin ring re-appeared at a new location (Figures 6B and 6C).

**Figure 6.**
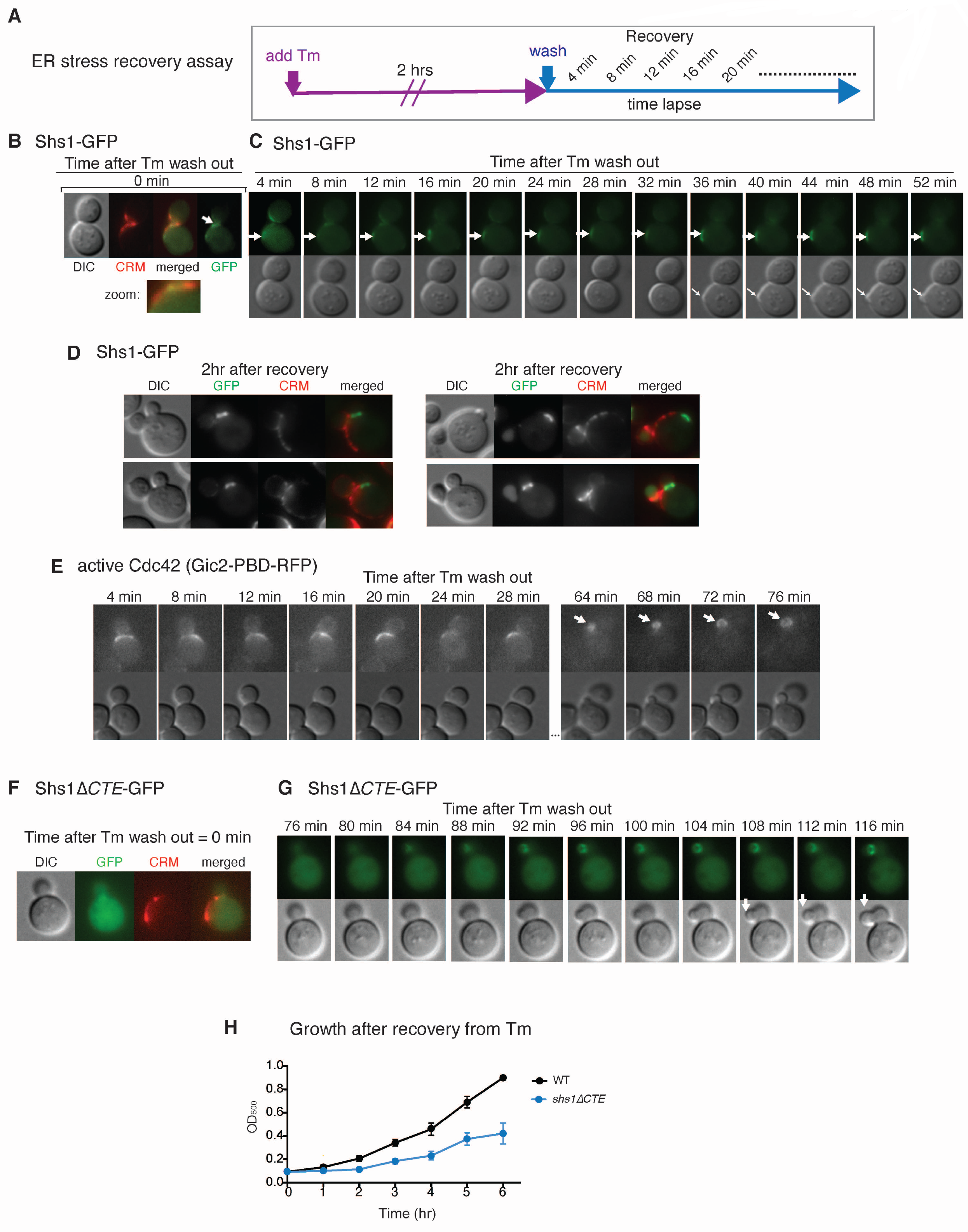
Septin ring translocation to CRMs regulated by the CTE of Shs1 dictates re-entry to the cell cycle when ER stress is recovered. (A) Experimental design for the ER stress recovery assay. Cells carrying Shs1-GFP were treated with 1µg/ml Tm for 2 hrs. Then, Tm was removed by washing the cells with SC medium without Tm. After, cells were monitored by live-cell imaging. In this assay, CRMs were visualized by staining with WGA-594.
(B, C) Recovery assay for WT cells expressing Shs1-GFP according to (A) to follow septin dynamics (Shs1-GFP) and bud emergence. The first frame of the time-lapse (time 0) is shown in (B) along with CRM staining. Subsequent frames are shown in (C). White arrow indicates bud emergence. See also Movie S6.
(D) Location of Shs1-GFP and CRMs in WT cells upon ER stress recovery for 2 hrs. As cells re-enter the cell cycle, a new bud emerges and Shs1-GFP localizes to the new bud. The initial bud was not re-used even after cells recovered from ER stress.
(E) Recovery assay to monitor active Cdc42 (Gic2-PDB-RFP) in WT cells according to (A). Active Cdc42 started to appear within 4 min after Tm wash and a new bud started to emerge at ∼60 min after Tm wash. Note that active Cdc42 appeared at the cortex of the newly emerging daughter cell.
(F, G) Recovery assay for Shs1-ΔCTE-GFP, which remained dispersed immediately after Tm recovery. The first frame of the time-lapse (time 0) is shown in (F) along with CRM staining. Subsequent frames are shown in (G). White arrow indicates bud emergence. See also Movie S7.
(H) Growth of WT or shs1-ΔCTE cells after ER stress recovery. Mean and SE were calculated based on values from at least three independent experiments.

This was followed by the emergence of a new bud after ∼50 min of recovery time (Figure 6C, Movie S6). This result is in agreement with our previous report that the original daughter cell is never re-used when cells recover from stress (Babour et al., 2010). As the original daughter cell was not utilized, we observed two budded cells in the early phase of recovery; however, the septin ring was only localized to the bud neck of the newly emerged daughter cell (Figure 6D).

Furthermore, during recovery, activated Cdc42 was polarized to the new presumptive bud site (Figure 6E). These results are consistent with our observation that the previously translocated septin ring moved from CRMs to a new incipient bud site, allowing cells to re-enter the cell cycle. As such, we most frequently observed new daughters emerged from sites adjacent to CRMs in these recovery assays.

In contrast to WT cells, both the re-appearance of the septin ring and bud emergence in Shs1Δ*CTE* cells were significantly delayed after recovery from ER stress (Figures 6F and 6G, Movie S7). The slower kinetics of the re-appearance of the septin ring after Tm removal and the new bud emergence was further reflected in the slower growth during the early recovery phase of 6 hrs (Figure 6H). Another striking difference with *shs1ΔCTE* cells was the specific location of the bud emergence: the new daughter cell emerged from within the existing daughter cell, instead of emerging from the mother cell as in WT (Figures 6B and 6G).

We further tested the importance of septin ring movement to CRMs in *bem1Δ* cells, in which the septin ring was transferred outside of the bud neck in a discrete location outside of CRMs under ER stress (Figures 5G and S6A, Movies S8). Interestingly, during the recovery process, the septin ring did not move from this transferred location in response to ER stress. In addition, the new bud emerged at this same location, although the kinetics of the bud emergence was very slow (Figures S6B and S6C). Thus, these results unveiled that the transfer of the septin ring from the bud neck specifically to CRMs is important for timely recovery and re-entry into the cell cycle.

### The impact of aging on ERSU cells

Establishment of a negative polarity cue (or disruption of a polarity cue) by ER stress-induced septin ring translocation to CRMs raises an interesting question regarding aged cells with multiple CRMs. After each cell division, bud scars accumulate on the cortex of the cell. Thus, aged cells have accumulated multiple CRMs (Powell et al., 2003; Sinclair et al., 1998). Taking this into account, how might aged cells respond to ER stress-induced septin ring translocation? Among asynchronously growing cells, over 50% are naive cells with only the birth scar (we refer to these as *‘young cells’*). Here, we examined how ER stress impacts cells that have more than three CRMs (we refer to these as *‘aged cells’*). Specifically, we investigated septin ring translocation in aged cells in response to ER stress. First, we quantified the levels of Nba1 at each bud scar in aged cells (Figures 7A and 7B).

**Figure 7.**
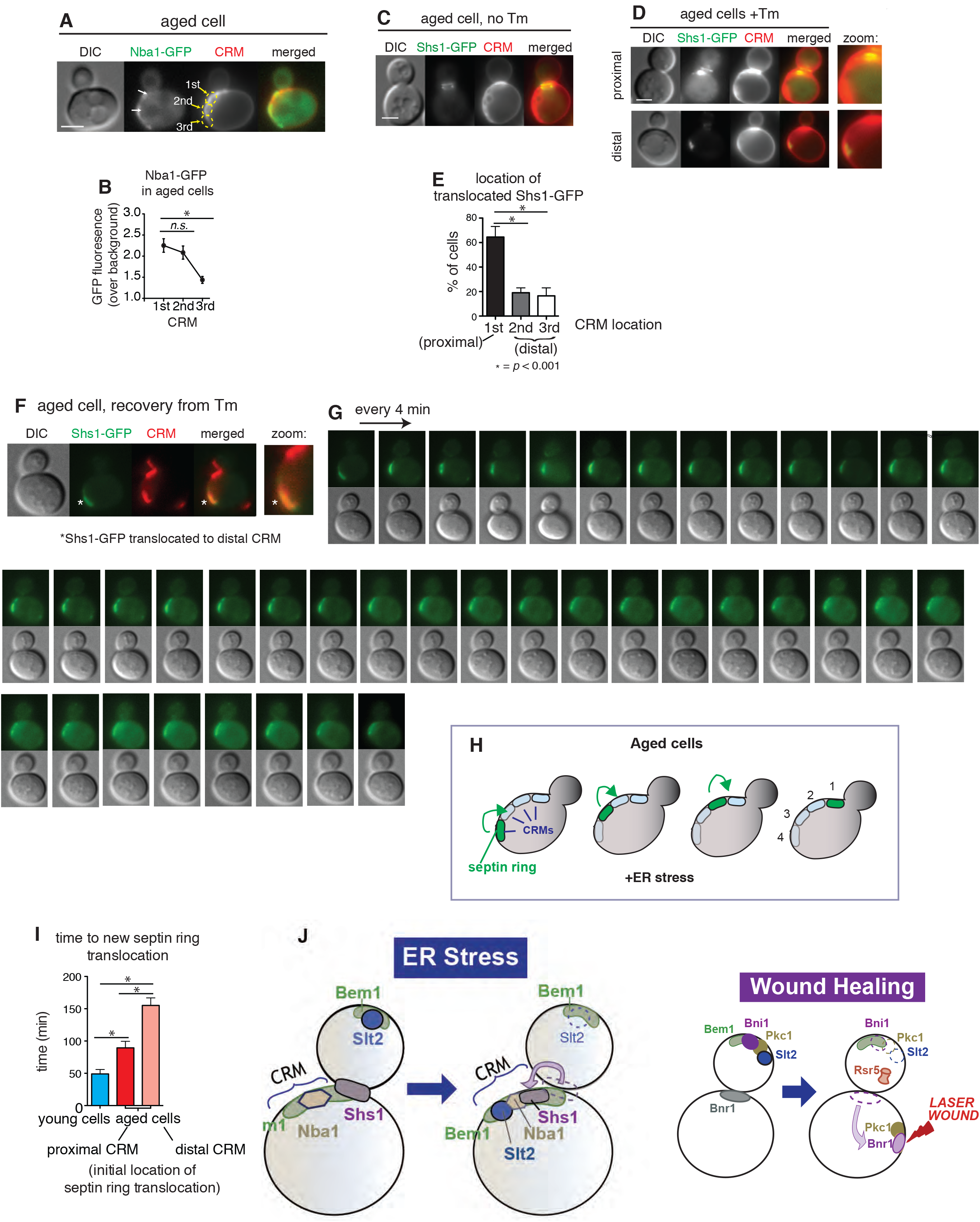
Septin ring behaviors in aged ER stressed yeast cells with multiple bud scars. (A, B) Nba1-GFP localization in Tm-treated (1 µg/ml) ‘*aged’* cells (A) and quantification of Nba1-GFP signals at different CRMs (B). Note that ‘*aged c*ells’ refers to cells two or more CRMs. “1^st^” “2^nd^” and “3^rd^” refer to the position of the CRM relative to the bud neck. *p* values comparing differences in GFP levels are indicated as: ns, not significant; *, p<0.001.
(C-E) Shs1-GFP localization in aged cells that were (C) untreated or (D) treated with Tm. Quantification of Shs1-GFP localization at each CRM is shown in (E). “Proximal” and “distal” refers to the location of the CRM relative to the bud neck. *p* values comparing differences in % of cells are indicated as *, p<0.001.
(F) ER stress recovery assay for *‘aged cells’* with Shs1-GFP transferred at the distant location immediately after removal of Tm.
(G, H) Time lapse of *‘aged cells’* with Shs1-GFP transferred at the distant location every 4 min after the removal of Tm and washing of cells (G). Graphic representation of septin ring localization during the time lapse experiment (H).
(I) Average time required for the reappearance of the septin ring at the new bud site, which was indicative of re-entry into the cell cycle, after ER stress recovery. *p* values were calculated to compare young cells and aged cells, and to compare aged cells with septin ring at either proximal or distal CRM prior to recovery in % of cells. *, p<0.001.
(J) In order for cells to effectively survive ‘*ER stress’*, a cellular strategy of hijacking of components that support cell growth and temporarily re-assembling them at a different location as an emergency complex allows proper handling of ER stress. Specifically, mobilizing components involved in the polarized growth at the bud tip, such as Bem1, Cdc24, and Slt2 and their re-assembly at CRMs along with association with the septin ring will achieve cell cycle halting while re-establishment of the ER functions is handled. Mobilization and re-formation of the polarized cell growth components may be a general emergency strategy for cells to cope with different types of stress or catastrophe. In response to a laser-induced wound, for example, components involved in polarized growth, such as Bni1 and Pkc1 (an upstream kinase of Slt2 MAP kinase), become mobilized from the bud tip, and their re-localization at the wound site allows for wound healing while cell growth is temporarily halted (Kono et al., 2012).

Although Nba1-GFP was present at each bud scar, it was highest in the one most proximal to the bud neck (Figure 7B), consistent with a previous report (Meitinger et al., 2014). The yeast strain used for these experiments has a BY genetic background; therefore, a new daughter cell should form axially. Based on our finding that Nba1 is a binding partner of Shs1 at CRMs, we tested if the septin ring transferred to the most recent CRMs during ER stress. In aged cells grown under normal growth conditions, the septin ring remained localized at the bud neck (Figure 7C). Upon ER stress induction, the septin ring transferred frequently (∼60%) to the bud scar most proximal to the bud neck, and less frequently (∼20%) to the more distal bud neck (Figures 5C-5E), indicating that septin rings translocate based on Nba1 levels.

### Age impacts the ability to undergo ER stress recovery

We next examined the effect of translocation of septin rings to older bud scars with less Nba1 (distal bud scars). Using the recovery assay described above (Figure 7A-7E), we monitored ER-stressed and aged cells with septin ring translocated to the distal bud scar (Figures 7F and 7G). Upon removal of the ER stress-inducing drug, the translocated septin ring moved away from the distal bud scar and re-localized to the more recent (newer) bud scar (Figure 7F-7I). Ultimately, septin rings ended up at the most proximal site to the bud neck through stepwise re-localization of bud scars in the middle. The amount of time required for aged cells to translocate the septin ring at the distal CRM to the most recent (proximal) CRM was significantly delayed when compared to naïve cells (Figures 7F-7H). Strikingly, despite the translocation of septin rings, we did not observe the emergence of a new daughter cell even after 120 min of ER stress recovery (Figure 7G). This is in stark contrast to naïve *WT* cells, where we detected re-entry into the cell cycle, scored by new bud emergence, within 90 min. Importantly, the recovery kinetics for aged WT cells was even slower than for *shs1ΔCTE* cells (Figure 6G). Thus, while ER stress-induced re-localization of septin ring could occur, the presence of multiple bud scars in aged cells diminished the ability of the cells to re-enter the cell cycle.

## DISCUSSION

One of the most important elements of cell survival is the correct segregation of cellular contents during cell division. Previously, we reported that the ERSU pathway, a cell cycle checkpoint, ensures the inheritance of functional ER during the cell cycle (Babour et al., 2010; Pina et al., 2018; Pina et al., 2016; Pina and Niwa, 2015). When the ER is stressed, the ERSU pathway blocks the inheritance of stressed ER and cytokinesis, thereby preventing generation of cells lacking sufficient levels of functional ER. Our previous work uncovered that cytokinesis block occurs by mislocalization of the septin ring, a critical component of cytokinesis; however, the mechanism of this mislocalization and its potential role beyond cytokinesis block remained elusive. Here, we found that this movement of septin rings to CRMs occurs by partially overriding the Nba1-dependent negative polarity establishment component. In addition, we found that Nba1 allows Bem1 at CRMs to find Cdc24 and Slt2; thus, CRMs serve as sites to gather key components in anticipation of cell cycle re-entry. Under normal growth, Nba1 at CRMs blocks both septin ring formation and Cdc24 association to CRM-localized Bem1, leading to the block of Cdc42 activation for polarized growth(Meitinger et al., 2014). Under ER stress, Nba1 is required for septin ring transfer to CRMs (Figure 7I). Furthermore, Nba1 allows Cdc24 to associate with Bem1, while still preventing Cdc42 from being activated. Bem1 localized at CRMs serves as the only binding site for activated Slt2, an ERSU MAP kinase. Taken together, our findings suggest that previous cell division sites, CRMs, serve as a reservoir for gathering components that aid in *‘cell cycle re-entry’* following recovery from ER.

In addition to the well-known role of septins in polarized growth and cell division (Hartwell, 1974), recent studies have shown that septins are involved in other aspects of cell polarity, including 1) the formation of cytoskeletal scaffolding structures (Mostowy and Cossart, 2012); 2) the formation of a diffusion barrier in the plasma membrane and ER (Caudron and Barral, 2009); and 3) lending definition to daughter-cell differentiation during bud emergence (Okada et al., 2013). All of these functions require septin rings to act as relatively static and passive landmarks during most of the cell cycle. The lack of septin ring formation at the bud neck prior to budding should trigger the Swe1-dependent morphogenesis check point or mitotic delay (Barral et al., 1999). This highlights the importance of the timely placement and generation of the septin ring at the site of polarized growth for initiating a new round of the cell cycle. By contrast, an unprecedented translocation of septin rings to CRMs enables cells to retain cellular abilities to re-enter cell cycle during the ER stress.

Based on time-lapse experiments, septin ring translocation to CRMs appears to occur via a transfer of a septin ring complex that is initially formed at the bud neck. In response to ER stress, we noticed that the diameter of the septin ring complex appeared to become smaller than the septin ring at the bud neck prior to ER stress. This suggests that ER stress causes a septin ring to take a more compact structure. Alternatively, some of the septin subunits initially formed at the bud neck may fall off to generate different septin rings at CRMs. At this point, we cannot rule out the possibility that some of the septin subunits disassemble at the bud neck and reassemble to a ring at CRMs. Supporting such a mechanism, a recent study showed that septin ring subunits gradually transfer from the previous bud neck to the incipient bud site (Chen et al., 2011). Regardless of the mechanism, the most important factor that distinguishes ER stress-induced septin ring is its transfer to CRMs rather than the incipient bud site.

The localization of septin rings at CRMs is also accompanied with other unexpected changes. Under normal growth, septin ring formation is coordinated with polarized cell growth. Under ER stress, the presence of septin rings at CRMs was not associated with bud emergence or polarized bud growth. Furthermore, under normal growth conditions, yeast cells have an elaborate mechanism involving a Cdc42 antagonist, Nba1, that prevents polarized growth from CRMs by interfering with the Cdc24 association with Bem1 localized at CRMs. Surprisingly, we found that Nba1 remained localized at CRMs under ER stress, but underwent certain character changes while retaining other features. For example, septin ring formation is normally coordinated with Cdc42 activation and polarized bud growth, but these events did not take place in septin rings transferred at CRMs under ER stress. Further, both Cdc24 and Slt2, but not Cdc42, localize to CRMs upon binding to Bem1. These results reveal that ER stress incapacitates a part of Nba1 functions, such that septin ring formation and Cdc24 localization can take place while Cdc42 activation or initiating polarized growth are continued to be blocked.

How can Bem1-Cdc24 be localized at CRMs in the presence of Nba1? Studies show that Bem1 serves two main functions: 1) as a scaffolding protein for a Cdc42-regulating effector, such as Cdc24 and 2) as a scaffolding protein for a Cdc42 effector p21-activated kinase (PAK), such as Cla4 or Ste20 (Atkins et al., 2008; Kozubowski et al., 2008). PAK phosphorylates Cdc42 while polarized growth is being established in yeast cells. This process helps ensure that the temporal and spatial regulation of Cdc42 activity determines the site and timing of symmetry breaking within the cell surface. Even after bud emergence, Bem1 remains at the bud tip until the direction of bud growth switches from an orthogonal to a bilateral direction. Our finding that Bem1 interacts with Slt2 during ER stress suggests that Slt2 may mediate one of the molecular switches that disassembles the Bem1-Cdc24 complex from the bud tip and assembles it at CRMs. Interestingly, Slt2 appears to be temporally activated during the cell cycle at around the time when the polarized growth switches directions even in the absence of ER stress (Li et al., 2010). Thus, under normal growth, the Slt2-Bem1 association may ultimately dictate Cdc42 activity in a spatially and temporally regulated manner to establish a switch of polarized growth. Upon ER stress activation, the Slt2-Bem1 interaction occurred at CRMs, which, as a consequence, may generate “hyper-activated” Bem1 or partially weaken Nba1 function. Thus, this process somehow facilitates Bem1-Cdc24 to localize to CRMs even in the presence of Nba1. However, the continued Nba1 presence may contribute to disconnecting the activation of Cdc42. Furthermore, the interaction of Bem1, Cdc24, and activated Slt2 might allow septin rings to translocate to CRMs even in the presence of a negative regulator such as Nba1. In fact, in *bem1Δ* cells, the septin ring (Shs1) ends up at the incipient bud site rather than at CRMs in response to ER stress, whereas in *nba1Δ cells,* some portions of the septin ring still transferred to CRMs. These observations are consistent with the idea that septin ring translocation to CRMs also requires Bem1 (Figure S7).

Importantly, the lack of gathering of ‘*cell cycle re-entry’* inducers at CRMs in ER-stressed *bem1Δ* or *shs1-ΔCTE* cells underscores the functional significance of concentrating Slt2, Cdc24, and septin rings at CRMs under ER stress. In both cases, the loss of transferred septin ring at CRMs significantly delays the cell’s ability to re-enter the cell cycle even when ER function is re-established. Thus, ER stress-induced septin ring transfer to CRMs represents a key event in anticipation of reestablished cellular competence of resuming cell cycle division following ER stress recovery.

Interestingly, upon re-entry into the cell cycle, we found that the original daughter cell was never utilized. Instead, a new bud emerged from the original mother cell. Under normal growth, establishing polarized growth requires an intrinsic competition between different foci of activated Cdc42 and its upstream GEFs and GAPs, which are aided by positive feedback to establish the “winning” foci for polarized growth that leads to the bud’s emergence from that foci (Wu et al., 2015). Once ER functional homeostasis is re-established, polarized growth can be re-established by re-mobilizing these components to the outside of the inhibitory zone that is defined by CRMs. Indeed, we found that upon recovery from ER stress new buds emerged from the sites directly adjacent to CRMs, retaining the usual axial budding pattern seen in haploid yeasts. The kinetics of the cell cycle recovery was significantly diminished in both *shs1-ΔCTE* and *bem1Δ* cells, in which the septin rings, Bem1, Cdc24, and Slt2 failed to gather at CRMs upon ER stress. Thus, these findings underscore the functional significance of strategically localizing key components at CRMs for resuming polarized growth in order to re-enter the cell cycle.

An interesting implication associated with CRMs as the site of septin ring translocation is aging, as cells accumulate CRMs as they undergo replication cycles. Therefore, unlike young cells, aging mother cells with multiple CRMs provide more choices for septin rings and other key components to congregate. Although young and aged cells appear to be equally effective at translocating septin rings, older cells with multiple CRMs struggle at re-localizing septin rings during recovery. Nba1 is present in several CRMs in such older cells. In addition, as cells age further, some of the CRMs may not have Nba1, which could alter the nature of septin ring transfer to occur at CRMs. One of the challenges of septin ring transfer to CRMs with decreased levels of Nba1 or little Nba1 includes strategies to block polarized growth.

Interestingly, UPR activation is also significantly slower in older cells (data not shown). Whatever the mechanisms might be, the prolonged kinetics of ER stress and UPR induction may provide additional time to gather components key to cell cycle re-entry. We found that septin rings transferred to older CRMs and then kept moving between CRMs until they reached the most recent CRMs before re-initiating the cell cycle. This might reveal that Nba1 concentration dictates the establishment towards recovery state. Furthermore, strong evidence suggests that during replicative aging, aging factors such as extra-chromosomal DNA circles and damaged-protein aggregates that decrease fitness of aged cells accumulate asymmetrically in the aging mothers (Erjavec et al., 2007; Higuchi-Sanabria et al., 2014; Shcheprova et al., 2008; Singh et al., 2017; Spokoini et al., 2012); this may also contribute to the delay of re-entry into the cell cycle during ER stress.

There is a precedent for a molecular strategy to respond to cellular emergencies by re-directing polarized cell growth components. For example, the mobilization/re-organization of Bni1—which regulates polarized cytoskeleton and Pkc1, an upstream kinase of Slt2—plays a major role in recovery from a laser-induced wound on the cell surface of yeast (Kono et al., 2012) (Figure 7I). The driving force behind halting polarized growth and mobilizing the necessary components for cell membrane surface repair is generated from rapidly degrading Bni1, a formin that nucleates actin filaments, via proteasomes. Concomitantly, Bnr1, another formin is mobilized to reach to the wound site to generate s ‘*wound recovery complex*’ along with Pkc1. A conceptual parallel can be found in ER-stressed cells: in response to ER stress, re-localization of Bem1, Slt2 kinase, and Cdc24 from the bud tip to CRMs is coordinated with the timely induction of the ERSU pathway. Re-establishment of the polarized growth and a platform for inheritance of the ER in the emerging daughter cell can be facilitated by the presence of this *‘cell cycle recovery complex’* at a reservoir of regulators of polarized growth at CRMs. Thus, our ER stress studies and those examining laser-induced wounds may have revealed an underlying principle and cellular strategy of handling catastrophes: by linking cell growth with the handling of a specific cellular stress, cells mobilizing components involved in polarized cell growth and re-functions them to take care of a specific cellular stress. Such a strategy ensures a break on the continued growth and provides effective means to handle stress recovery. One important element of such a stress-handling strategy is to retain the ability to resume polarized growth. CRMs may provide an ideal location for transferring ‘*cell cycle re-entry’* components under ER stress in the absence of specific targets such as wounded sites in the cell.

Finally, as ER stress is conserved among eukaryotic cells, it is tempting to speculate that a similar mechanism might exist for effectively handling ER stress in mammalian cells (Chen et al., 2013; Ettinger et al., 2011; Kuo et al., 2011; Pohl and Jentsch, 2009; Thieleke-Matos et al., 2017). While the molecular basis of the CRM and its constituents are unique to yeast cells, recent studies have suggested that the mid body (MB) formed at the cleavage furrow during cytokinesis may be a functional equivalent of CRMs. Structurally, the MB takes a ring-like shape that resembles CRMs. As CRMs represent prior cytokinesis sites, the MB is also formed during cytokinesis and plays a role in cell division. While the exact MB constituents differ from those of CRMs, recent studies revealed that the MB also retains the post-mitotic structure. Specifically, upon division of mammalian two daughter cells, the MB ends up in one of the daughter cells. Ultimately, many MBs can be removed from the cell by a few mechanisms including autophagy, although its half-life appears to differ depending on the cell type. The half-life of an MB appears to dictate the pluripotency of stem cells. For example, stem cells with a long-lived or persistent MB normally retain pluripotency. By contrast, during asymmetric division of stem cells, in which one cell retains pluripotency and the other differentiates into a specific cell type, the extent of potency is correlated with the half-life of the MB. Given our findings on CRMs, it will be interesting to test if pluripotency of MBs change in response to ER stress.

## Inventory

### Supplementary Figures and Figure Legends

**Figure S1.**
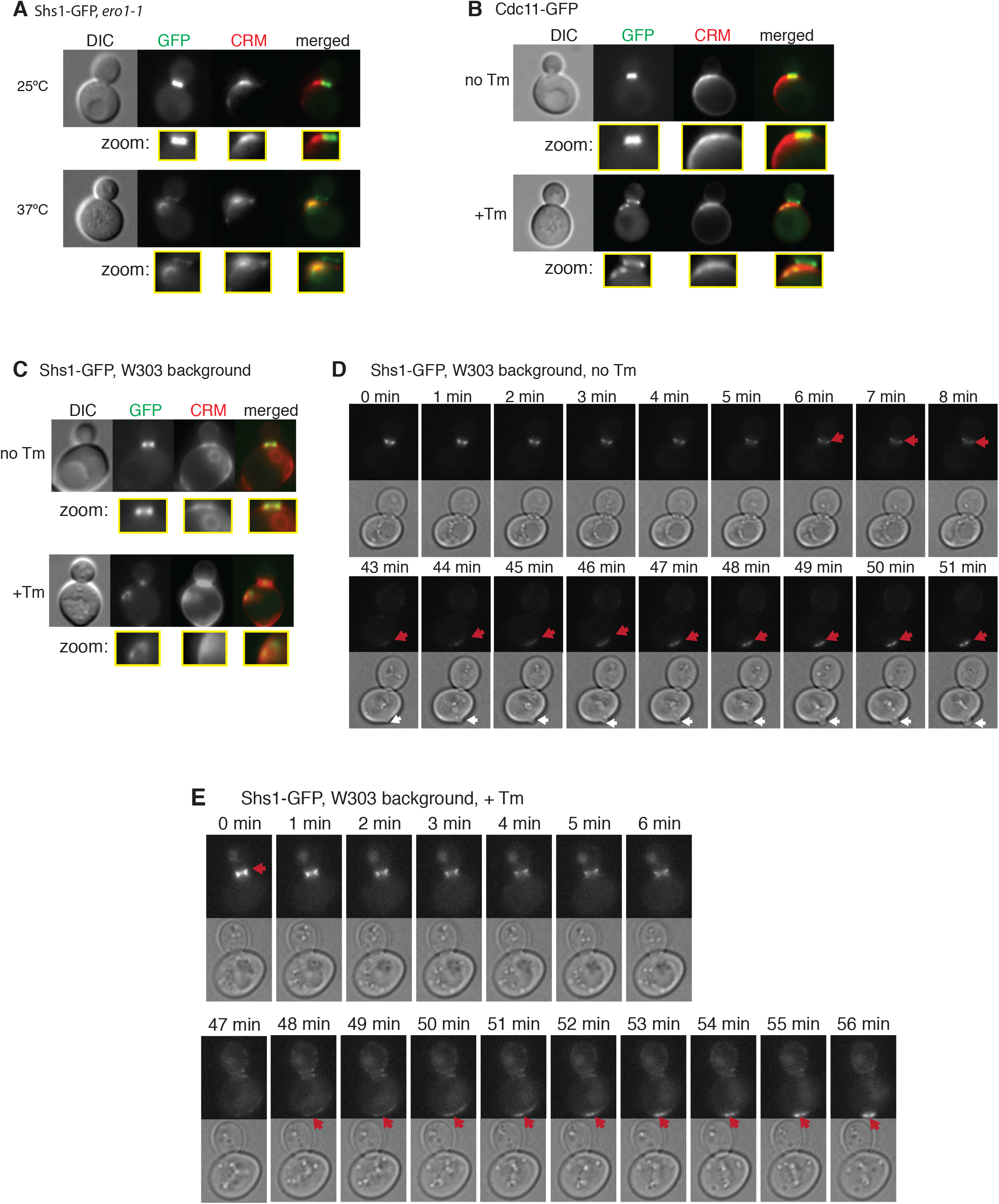
Septin rings are mislocalized at CRMs regardless of ER stress types and of budding pattern differences. (A) Shs1-GFP expressed in *ero1-1* cells grown at permissive (25**°**C) and non-permissive (37**°**C) temperatures. In all panels, zoomed images show cropped images from the bud neck regions, and CRMs were visualized by CWS.
(B) Cdc11-GFP in WT cells grown without or with 1µg/ml of Tm.
(C) Shs1-GFP in WT cells of the W303 background grown without or with 1µg/ml of Tm.
(D and E) Time-lapse analysis of Shs1-GFP translocation in W303 cells either untreated (no Tm) (D) or treated with 1µg/ml Tm (+Tm) (E).

**Figure S2.**
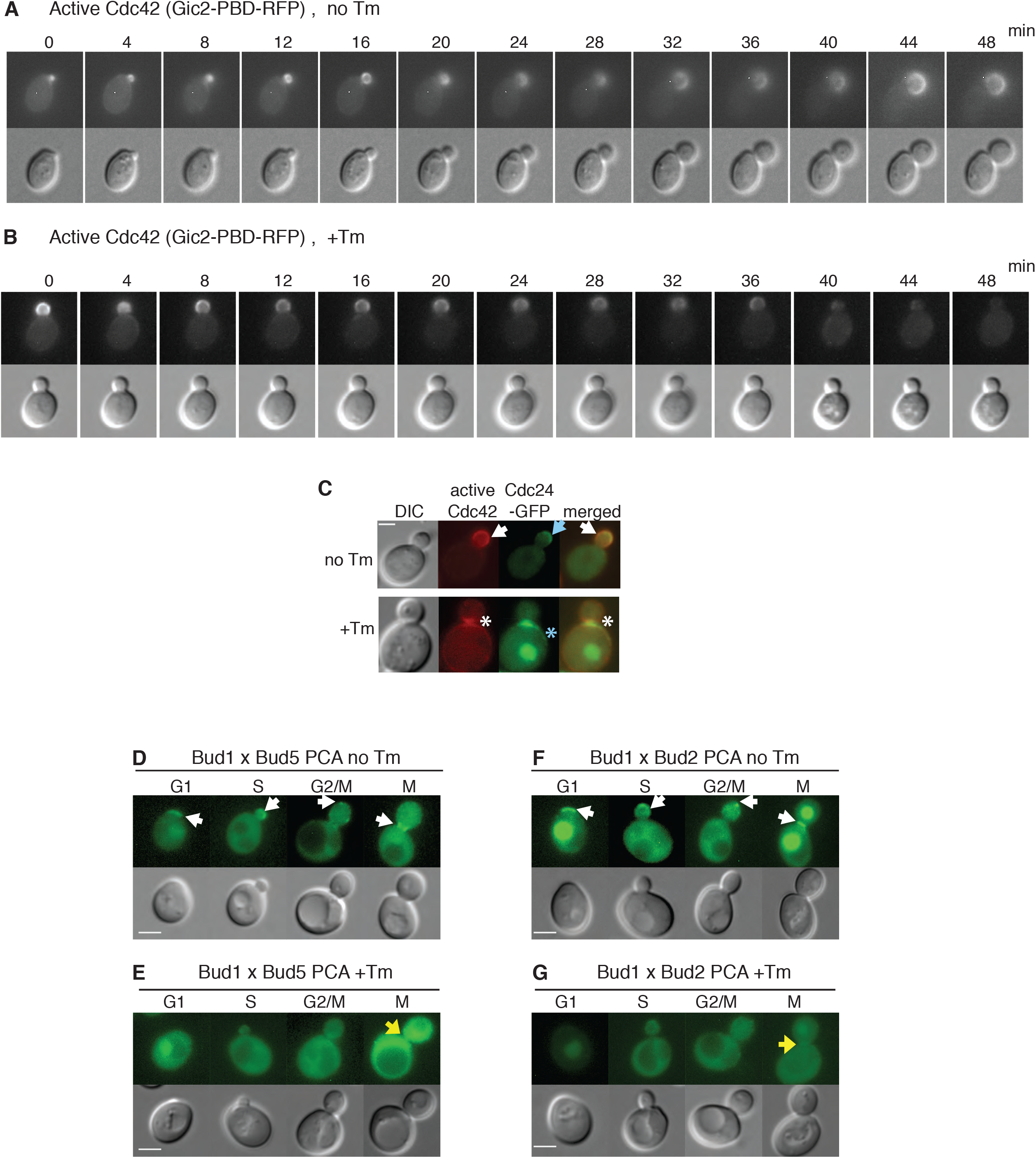
Cdc42 is inactivated in response to ER stress induction. (A and B) Time-lapse analysis of active Cdc42 (Gic2-PBD-RFP) in WT cells with or without 1µg/ml Tm in microfluidics chambers.
(C) Cells co-expressing Gic2-PBD-RFP and Cdc24-GFP were grown with or without 1µg/ml Tm. White arrows show Gic2-PBD-RFP, blue arrows show Cdc24-GFP, and asterisks show mislocalized Gic2-PBD-RFP (white) or Cdc24 (blue). Scale bar, 2 µm.
(D and E) Split-YFP PCA between Bud1 and Bud5. Cells were grown with or without 1 µg/ml Tm. White arrows show where Bud1 and Bud5 interacted in untreated cells (D) and a yellow arrowhead in (E) shows a small amount of Bud1 interaction with Bud5 in ER stressed cells.
(F and G) Split-YFP PCA between Bud1 and Bud2. Cells were grown without (F) or with 1 µg/ml Tm (G). Bud1 and Bud2 interacted in unstressed cells (white arrows), and very little interaction (yellow arrowhead) was detected during ER stress in an M phase cell.

**Figure S3.**
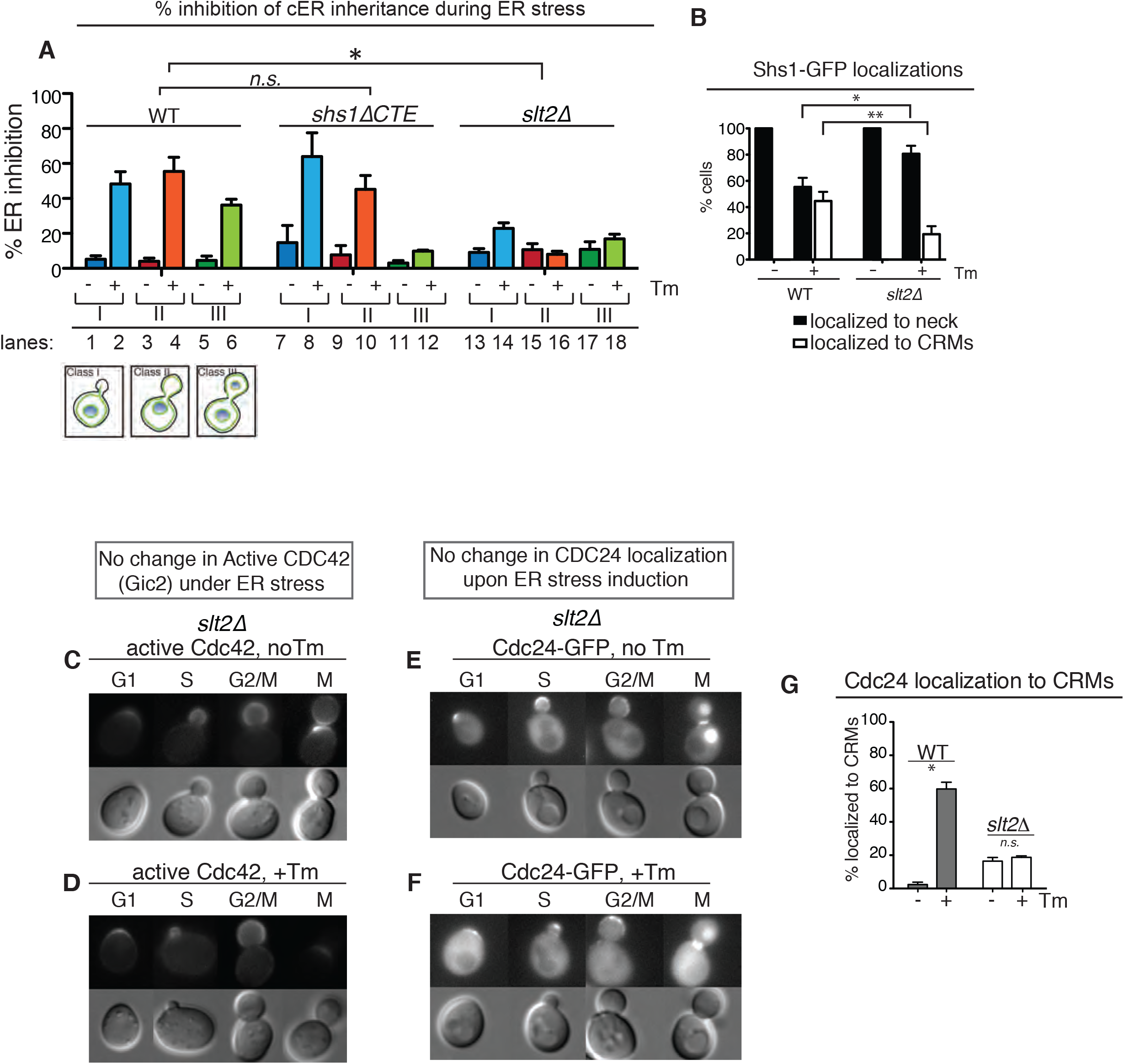
*slt2Δ* cells fail to induce the ERSU pathway and no change in Cdc24 localization upon ER stress induction. (A) The cER inheritance was blocked in ER-stressed *shs1ΔCTE* cells. The criteria for classes of cells used are as previously described (Babour et al., 2010): Class I represents cells with small buds of less than 2μm in diameter; Class II cells are cells with medium-large sized buds of 2 μm or larger; and class III are cells with large buds containing the inherited nucleus. Interestingly, the cER inheritance of ER-stressed class III *shs1ΔCTE* cells was not significantly affected. No significant level of the cER inheritance block occurred in ER-stressed *slt2Δ* cells. Two-way ANOVA comparing different strains’ responses to ER stress showed that *shs1ΔCTE* was not significantly different than WT (n.s.), but *slt2Δ* was (* represents *p* < 0.0001).
(B) Quantifications for Shs1-GFP localizations in WT and *slt2Δ* cells either untreated (-Tm) or treated with 1 µg/ml Tm (+Tm). *, *p* < 0.05; **, *p* < 0.01.
(C and D) Gic2-PBD-RFP in *slt2Δ* cells remained unchanged between untreated (C) and 1µg/ml Tm-treated (D) conditions.
(D and F) Cdc24-GFP in *slt2Δ* cells remained unchanged between untreated (D) and 1µg/ml Tm-treated (E) conditions.
(G) Quantification of Cdc24-GFP localized to CRMs in WT and *slt2Δ* cells with or without ER stress with Tm treatment (1µg/ml). T-tests comparing treated and untreated conditions showed that * = *p* < 0.001 and *n.s.* = not significant.

**Figure S4.**
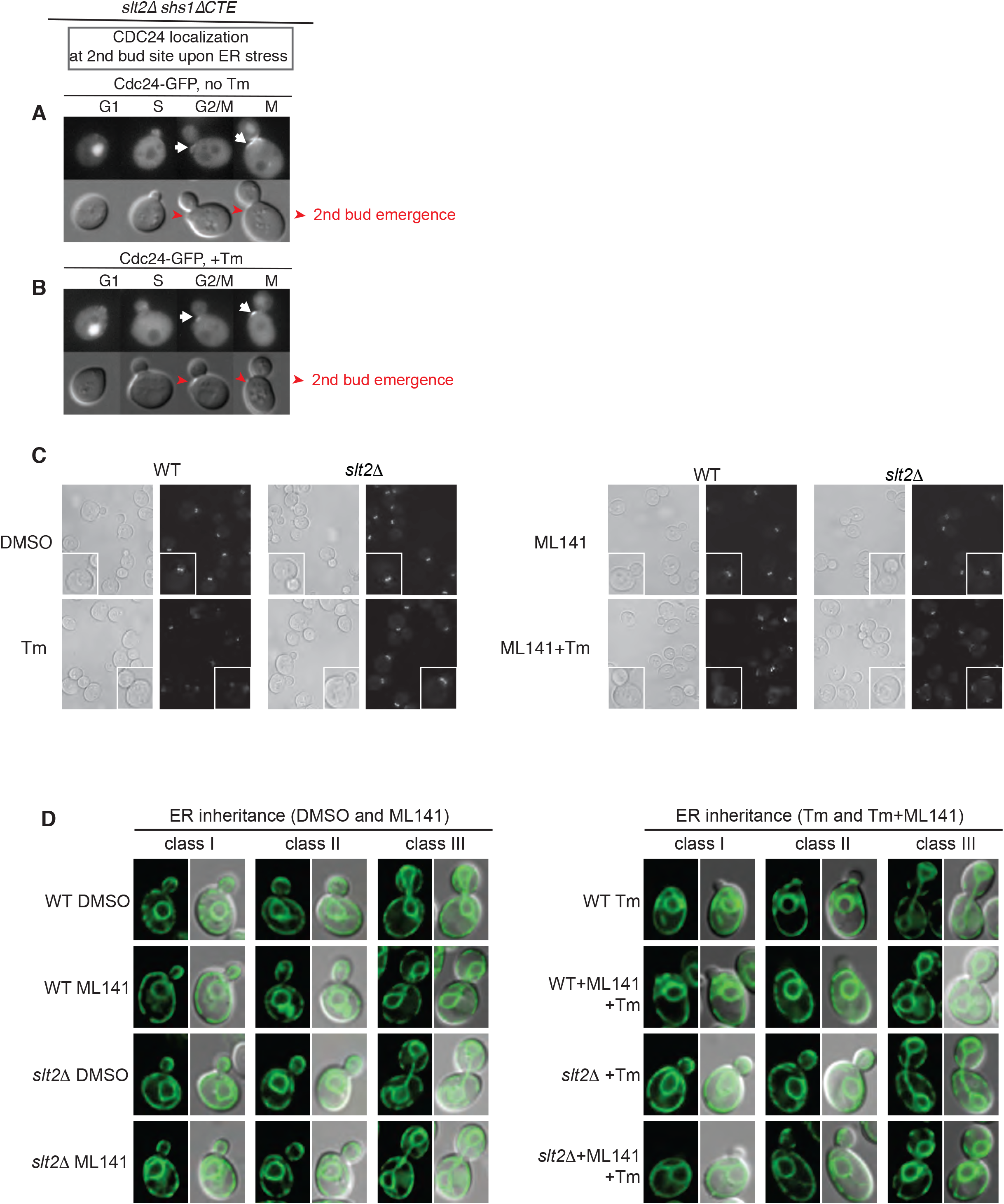
*slt2Δ shs1ΔCTE* cells fail to induce the ERSU pathway but can be rescued by a Cdc42 inhibitor. (A, B) Cdc24-GFP (white arrows) in *slt2Δ shs1ΔCTE cells* was localized to the incipient 2^nd^ bud site (indicated as red arrowheads) under both normal growth or upon ER stress by treatment with Tm.
(C,) Representative images of Shs1-GFP in WT and *slt2Δ* cells treated with DMSO, Tm (0.5 μg/ml), ML141 (20µM), or ML141 (20µM) plus Tm (0.5 μg/ml). Quantification of these experiments is shown in Figure 4H.
(D) Representative images of the cER inheritance in WT and *slt2Δ* cells, treated with DMSO, Tm (0.5 μg/ml), ML141 (20µM), or ML141 (20µM) plus Tm (0.5 μg/ml). Quantifications of these experiments are shown in Figure 4G.

**Figure S5.**
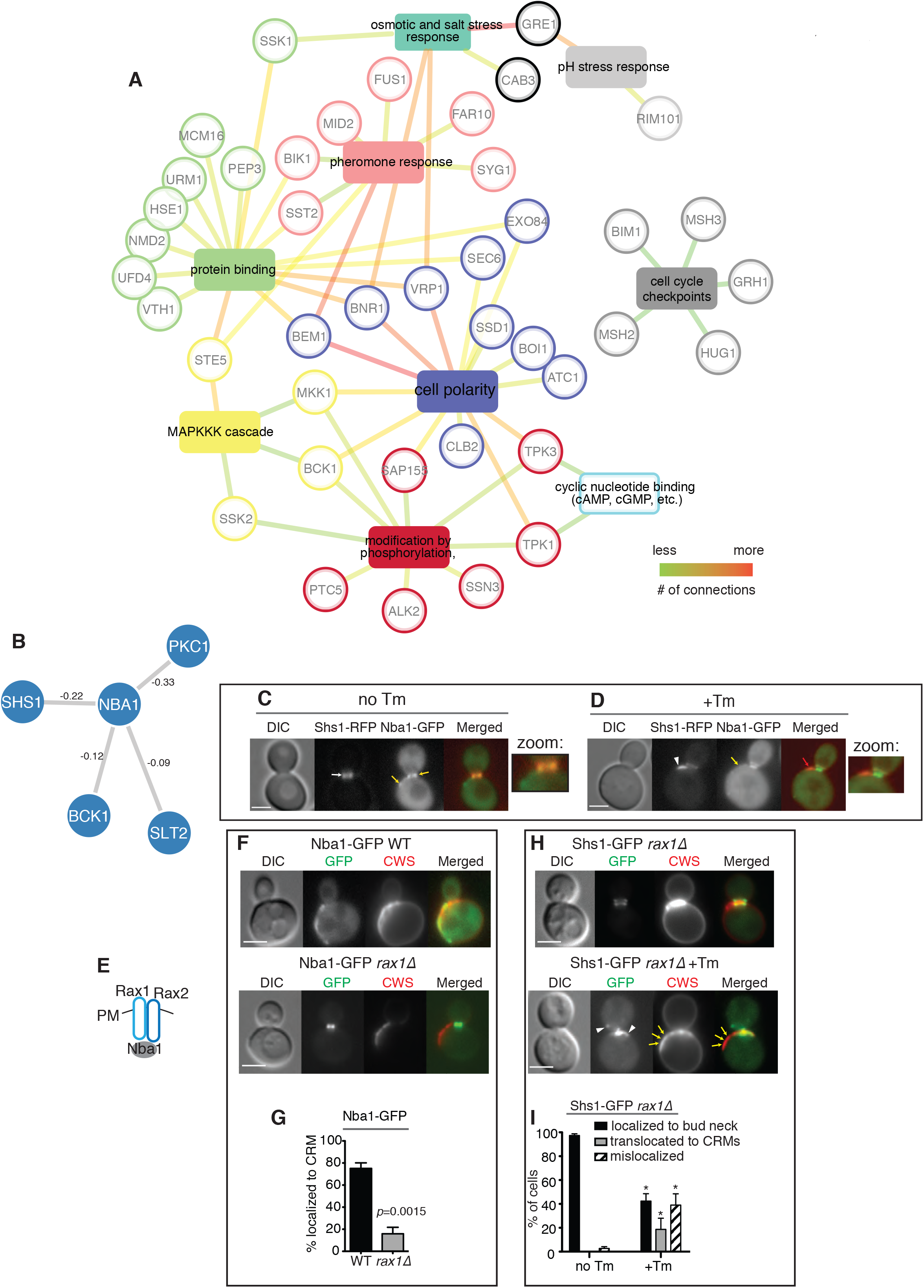
Identification of Bem1 as a Slt2-interacting protein. (A) Functional interaction map for Slt2 derived from protein-protein interactions identified in the split-DHFR screen. Nodes represent functional categories, and edges define associations with Slt2. Edge length and thickness indicate fold enrichment, which is defined as the number of proteins from the input dataset relative to the total number of proteins in a given category. For details on the analysis, see Materials and Methods. See also Table S1.
(B) Genetic interactions between *NBA1* and genes in the *SLT2* pathway as well as *SHS1* as identified by Costanzo et al. 2010. The strength of interactions are annotated on the edges.
(C D) Nba1-RFP colocalizes with Shs1-GFP at the bud neck in untreated cells (C) and at CRMs in Tm-treated cells (D).
(E) Rax1 and Rax2 are important for localization of Nba1 to the CRMs at the plasma membrane (PM) (Meitinger et al. 2014).
(F) Nba1-GFP localization in WT or *rax1Δ* cells. The loss of *RAX1* (*rax1Δ*) significantly diminisheD Nba1-GFP localization to CRMs and Nba1-GFP remained at the bud neck.
(G) Quantitation of Nba1-GFP localized at CRMs. T-test showed that Nba1-GFP localizations to CRMs was significantly reduced in *rax1Δ* compared to WT.
(H, I) Shs1-GFP localization in untreated and Tm-treated *rax1Δ* cells (H) and quantification of Shs1-GFP localization (I). Shs1-GFP localizes to the bud neck during normal growth, and it remained at the bud neck in ER-stressed *rax1Δ* cells. P values from t-tests comparing each of the Shs1-GFP localizations between no Tm and +Tm conditions is shown (* represents p < 0.001).

**Figure S6.**
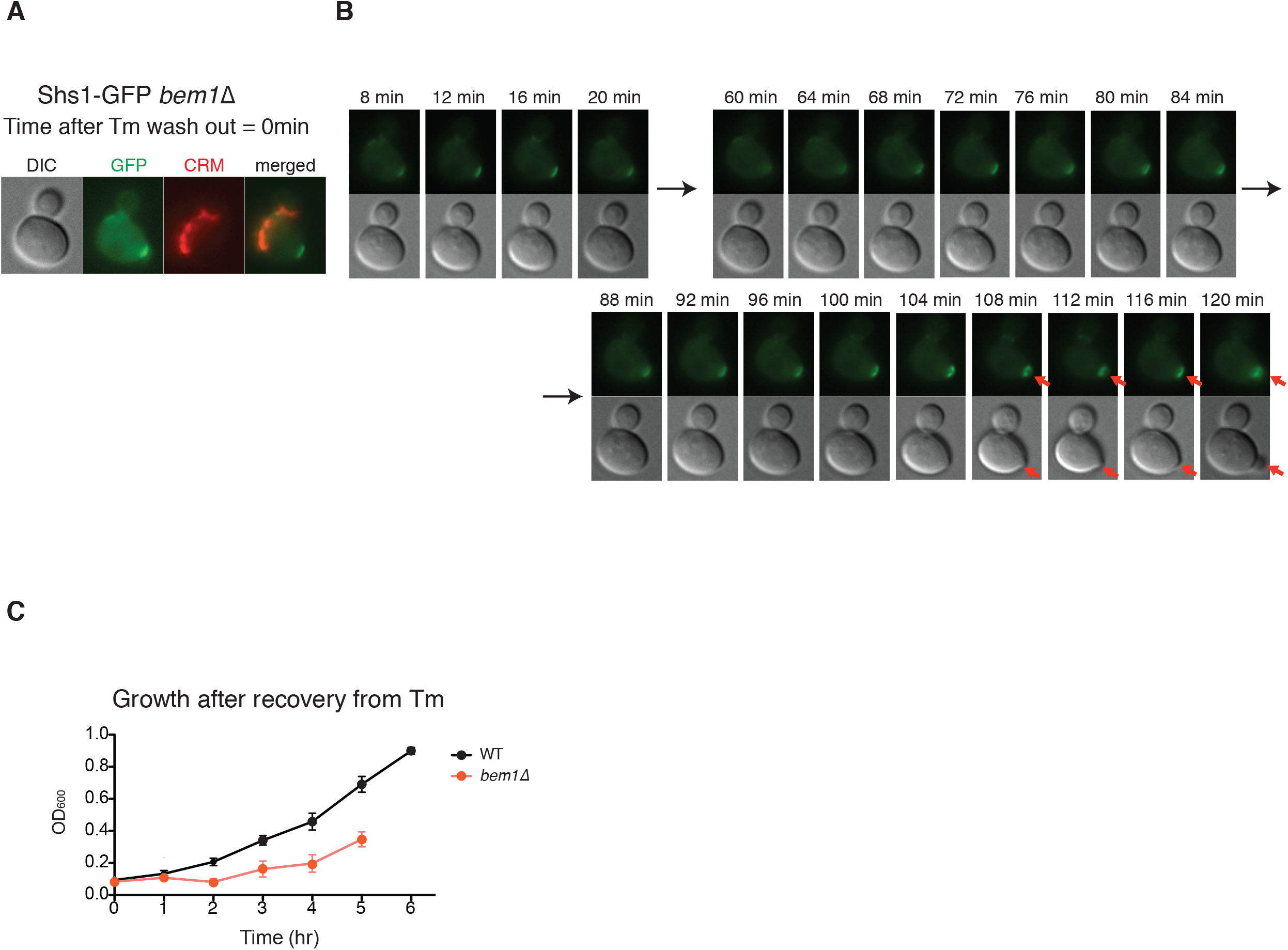
ER stress recovery of *bem1Δ* cells is diminished. (A, B) Dynamics of ER stress recovery of *bem1Δ* cells. ER stress recovery assay for *bem1Δ* cells was done according to Figure 6A. The first frame of the time-lapse (time 0) is shown in (A) along with CRM staining. Subsequent frames are shown in (B). Orange arrow indicates bud emergence after recovery. See also Movie S8.
(C) Cell growth after recovery from ER stress between WT and *bem1Δ* cells. Error bars, SE.

### Supplementary Movie Legends

Movie S1. Shs1-GFP translocation in an untreated WT cell. Related to Figure 1A.

Movie S2. Shs1-GFP translocation during ER stress in a WT cell. Related to Figure 1B.

Movie S3. Shs1-GFP translocation in an untreated WT cell of the W303 background. Related to Figure S1D.

Movie S4. Shs1-GFP translocation in W303 WT cell during ER stress. Related to Figure S1E.

Movie S5. Dynamics of Cdc42-GTP during ER stress. Related to Figure S2A.

Movie S6. Shs1-GFP dynamics in a WT cell that was first treated with Tm and then washed into normal medium for recovery. Related to Figure 6B.

Movie S7. Shs1ΔCTE-GFP dynamics during recovery from ER stress. Related to Figure 6G.

Movie S8. Shs1-GFP dynamics in a *bem1Δ* cell during recovery from ER stress. Related to Figure S6B.

### Supplemental movies

**Movie S1.** Shs1-GFP in WT cell untreated.

**Movie S2.** Shs1-GFP in WT cell +Tm.

**Movie S3.** Shs1-GFP in WT W303 cell, untreated.

**Movie S4.** Shs1-GFP in WT W303 cell +Tm.

**Movie S5.** Gic2-PBD-RFP in WT cell +Tm.

**Movie S6.** Shs1-GFP in WT cell recovering from ER stress.

**Movie S7.** Shs1Δ*CTE*-GFP cells recovering from ER stress.

**Movie S8.** Shs1-GFP in *bem1Δ* cells recovering from ER stress.

## References

1. Atkins, B.D., Yoshida, S., and Pellman, D. (2008). Symmetry breaking: scaffold plays matchmaker for polarity signaling proteins. Curr Biol 18, R1130–1132.

2. Babour, A., Bicknell, A.A., Tourtellotte, J., and Niwa, M. (2010). A surveillance pathway monitors the fitness of the endoplasmic reticulum to control its inheritance. Cell 142, 256–269.

3. Barlowe, C. (2010). ER sheets get roughed up. Cell 143, 665–666.

4. Barral, Y., Parra, M., Bidlingmaier, S., and Snyder, M. (1999). Nim1-related kinases coordinate cell cycle progression with the organization of the peripheral cytoskeleton in yeast. Genes Dev 13, 176–187.

5. Bechmann, L.P., Hannivoort, R.A., Gerken, G., Hotamisligil, G.S., Trauner, M., and Canbay, A. (2012). The interaction of hepatic lipid and glucose metabolism in liver diseases. J Hepatol 56, 952–964.

6. Bi, E., Maddox, P., Lew, D.J., Salmon, E.D., McMillan, J.N., Yeh, E., and Pringle, J.R. (1998). Involvement of an actomyosin contractile ring in Saccharomyces cerevisiae cytokinesis. J Cell Biol 142, 1301–1312.

7. Bos, J.L., Rehmann, H., and Wittinghofer, A. (2007). GEFs and GAPs: critical elements in the control of small G proteins. Cell 129, 865–877.

8. Caudron, F., and Barral, Y. (2009). Septins and the lateral compartmentalization of eukaryotic membranes. Dev Cell 16, 493–506.

9. Chen, C.T., Ettinger, A.W., Huttner, W.B., and Doxsey, S.J. (2013). Resurrecting remnants: the lives of post-mitotic midbodies. Trends Cell Biol 23, 118–128.

10. Chen, H., Howell, A.S., Robeson, A., and Lew, D.J. (2011). Dynamics of septin ring and collar formation in Saccharomyces cerevisiae. Biol Chem 392, 689–697.

11. Chen, R.E., and Thorner, J. (2007). Function and regulation in MAPK signaling pathways: lessons learned from the yeast Saccharomyces cerevisiae. Biochim Biophys Acta 1773, 1311–1340.

12. Chiou, J.G., Balasubramanian, M.K., and Lew, D.J. (2017). Cell Polarity in Yeast. Annu Rev Cell Dev Biol 33, 77–101.

13. Costanzo, M., Baryshnikova, A., Myers, C.L., Andrews, B., and Boone, C. (2011). Charting the genetic interaction map of a cell. Curr Opin Biotechnol 22, 66–74.

14. Denic, V., Quan, E.M., and Weissman, J.S. (2006). A luminal surveillance complex that selects misfolded glycoproteins for ER-associated degradation. Cell 126, 349–359.

15. Du, Y., Ferro-Novick, S., and Novick, P. (2004). Dynamics and inheritance of the endoplasmic reticulum. Journal of Cell Science 117, 2871–2878.

16. Erjavec, N., Larsson, L., Grantham, J., and Nystrom, T. (2007). Accelerated aging and failure to segregate damaged proteins in Sir2 mutants can be suppressed by overproducing the protein aggregation-remodeling factor Hsp104p. Genes Dev 21, 2410–2421.

17. Ettinger, A.W., Wilsch-Brauninger, M., Marzesco, A.M., Bickle, M., Lohmann, A., Maliga, Z., Karbanova, J., Corbeil, D., Hyman, A.A., and Huttner, W.B. (2011). Proliferating versus differentiating stem and cancer cells exhibit distinct midbody-release behaviour. Nat Commun 2, 503.

18. Fehrenbacher, K.L., Davis, D., Wu, M., Boldogh, I., and Pon, L.A. (2002). Endoplasmic reticulum dynamics, inheritance, and cytoskeletal interactions in budding yeast. Molecular Biology of the Cell 13, 854–865.

19. Feige, M.J., and Hendershot, L.M. (2011). Disulfide bonds in ER protein folding and homeostasis. Curr Opin Cell Biol 23, 167–175.

20. Field, C.M., and Kellogg, D. (1999). Septins: cytoskeletal polymers or signalling GTPases? Trends Cell Biol 9, 387–394.

21. Frakes, A.E., and Dillin, A. (2017). The UPRER: Sensor and Coordinator of Organismal Homeostasis. Mol Cell 66, 761–771.

22. Frand, A.R., and Kaiser, C.A. (1998). The ERO1 gene of yeast is required for oxidation of protein dithiols in the endoplasmic reticulum. Molecular Cell 1, 161–170.

23. Gladfelter, A.S., Pringle, J.R., and Lew, D.J. (2001). The septin cortex at the yeast mother-bud neck. Curr Opin Microbiol 4, 681–689.

24. Gustin, M.C., Albertyn, J., Alexander, M., and Davenport, K. (1998). MAP kinase pathways in the yeast Saccharomyces cerevisiae. Microbiol Mol Biol Rev 62, 1264–1300.

25. Hartwell, L.H. (1974). Saccharomyces cerevisiae cell cycle. Bacteriol Rev 38, 164–198.

26. Hereford, L.M., and Hartwell, L.H. (1974). Sequential gene function in the initiation of Saccharomyces cerevisiae DNA synthesis. J Mol Biol 84, 445–461.

27. Higuchi-Sanabria, R., Pernice, W.M., Vevea, J.D., Alessi Wolken, D.M., Boldogh, I.R., and Pon, L.A. (2014). Role of asymmetric cell division in lifespan control in Saccharomyces cerevisiae. FEMS Yeast Res 14, 1133–1146.

28. Kang, P.J., Sanson, A., Lee, B., and Park, H.O. (2001). A GDP/GTP exchange factor involved in linking a spatial landmark to cell polarity. Science 292, 1376–1378.

29. Kono, K., Saeki, Y., Yoshida, S., Tanaka, K., and Pellman, D. (2012). Proteasomal degradation resolves competition between cell polarization and cellular wound healing. Cell 150, 151–164.

30. Kozubowski, L., Saito, K., Johnson, J.M., Howell, A.S., Zyla, T.R., and Lew, D.J. (2008). Symmetry-breaking polarization driven by a Cdc42p GEF-PAK complex. Curr Biol 18, 1719–1726.

31. Kuo, S.C., and Lampen, J.O. (1974). Tunicamycin--an inhibitor of yeast glycoprotein synthesis. Biochem Biophys Res Commun 58, 287–295.

32. Kuo, T.C., Chen, C.T., Baron, D., Onder, T.T., Loewer, S., Almeida, S., Weismann, C.M., Xu, P., Houghton, J.M., Gao, F.B., et al. (2011). Midbody accumulation through evasion of autophagy contributes to cellular reprogramming and tumorigenicity. Nat Cell Biol 13, 1214–1223.

33. Li, X., Du, Y., Siegel, S., Ferro-Novick, S., and Novick, P. (2010). Activation of the mitogen-activated protein kinase, Slt2p, at bud tips blocks a late stage of endoplasmic reticulum inheritance in Saccharomyces cerevisiae. Mol Biol Cell 21, 1772–1782.

34. Lippincott, J., and Li, R. (1998). Dual function of Cyk2, a cdc15/PSTPIP family protein, in regulating actomyosin ring dynamics and septin distribution. J Cell Biol 143, 1947–1960.

35. Liu, D., and Novick, P. (2014). Bem1p contributes to secretory pathway polarization through a direct interaction with Exo70p. J Cell Biol 207, 59–72.

36. Lord, M., Laves, E., and Pollard, T.D. (2005). Cytokinesis depends on the motor domains of myosin-II in fission yeast but not in budding yeast. Mol Biol Cell 16, 5346–5355.

37. Madden, K., and Snyder, M. (1998). Cell polarity and morphogenesis in budding yeast. Annu Rev Microbiol 52, 687–744.

38. Marston, A.L., Chen, T., Yang, M.C., Belhumeur, P., and Chant, J. (2001). A localized GTPase exchange factor, Bud5, determines the orientation of division axes in yeast. Curr Biol 11, 803–807.

39. Meitinger, F., Khmelinskii, A., Morlot, S., Kurtulmus, B., Palani, S., Andres-Pons, A., Hub, B., Knop, M., Charvin, G., and Pereira, G. (2014). A memory system of negative polarity cues prevents replicative aging. Cell 159, 1056–1069.

40. Michnick, S.W., Ear, P.H., Manderson, E.N., Remy, I., and Stefan, E. (2007). Universal strategies in research and drug discovery based on protein-fragment complementation assays. Nat Rev Drug Discov 6, 569–582.

41. Mori, K. (2000). Tripartite management of unfolded proteins in the endoplasmic reticulum. Cell 101, 451–454.

42. Mostowy, S., and Cossart, P. (2012). Septins: the fourth component of the cytoskeleton. Nat Rev Mol Cell Biol 13, 183–194.

43. Nelson, S.A., Sanson, A.M., Park, H.O., and Cooper, J.A. (2012). A novel role for the GTPase-activating protein Bud2 in the spindle position checkpoint. PLoS One 7, e36127.

44. Oh, Y., and Bi, E. (2011). Septin structure and function in yeast and beyond. Trends Cell Biol 21, 141–148.

45. Okada, S., Leda, M., Hanna, J., Savage, N.S., Bi, E., and Goryachev, A.B. (2013). Daughter cell identity emerges from the interplay of Cdc42, septins, and exocytosis. Dev Cell 26, 148–161.

46. Onishi, M., Ko, N., Nishihama, R., and Pringle, J.R. (2013). Distinct roles of Rho1, Cdc42, and Cyk3 in septum formation and abscission during yeast cytokinesis. J Cell Biol 202, 311–329.

47. Park, H.O., Bi, E., Pringle, J.R., and Herskowitz, I. (1997). Two active states of the Ras-related Bud1/Rsr1 protein bind to different effectors to determine yeast cell polarity. Proc Natl Acad Sci U S A 94, 4463–4468.

48. Park, H.O., Kang, P.J., and Rachfal, A.W. (2002). Localization of the Rsr1/Bud1 GTPase involved in selection of a proper growth site in yeast. J Biol Chem 277, 26721–26724.

49. Pina, F., Yagisawa, F., Obara, K., Gregerson, J.D., Kihara, A., and Niwa, M. (2018). Sphingolipids activate the endoplasmic reticulum stress surveillance pathway. J Cell Biol.

50. Pina, F.J., Fleming, T., Pogliano, K., and Niwa, M. (2016). Reticulons Regulate the ER Inheritance Block during ER Stress. Dev Cell 37, 279–288.

51. Pina, F.J., and Niwa, M. (2015). The ER Stress Surveillance (ERSU) pathway regulates daughter cell ER protein aggregate inheritance. Elife 4.

52. Pohl, C., and Jentsch, S. (2009). Midbody ring disposal by autophagy is a post-abscission event of cytokinesis. Nat Cell Biol 11, 65–70.

53. Pollard, M.G., Travers, K.J., and Weissman, J.S. (1998). Ero1p: a novel and ubiquitous protein with an essential role in oxidative protein folding in the endoplasmic reticulum. Molecular Cell 1, 171–182.

54. Powell, C.D., Quain, D.E., and Smart, K.A. (2003). Chitin scar breaks in aged Saccharomyces cerevisiae. Microbiology 149, 3129–3137.

55. Pruyne, D., and Bretscher, A. (2000). Polarization of cell growth in yeast. I. Establishment and maintenance of polarity states. J Cell Sci 113 ( *Pt 3*), 365–375.

56. Pruyne, D., Legesse-Miller, A., Gao, L., Dong, Y., and Bretscher, A. (2004). Mechanisms of polarized growth and organelle segregation in yeast. Annual Review of Cell and Developmental Biology 20, 559–591.

57. Robinson, M.D., Grigull, J., Mohammad, N., and Hughes, T.R. (2002). FunSpec: a web-based cluster interpreter for yeast. BMC Bioinformatics 3, 35.

58. Ron, D., and Walter, P. (2007). Signal integration in the endoplasmic reticulum unfolded protein response. Nat Rev Mol Cell Biol 8, 519–529.

59. Rutkowski, D.T., and Kaufman, R.J. (2004). A trip to the ER: coping with stress. Trends Cell Biol 14, 20–28.

60. Shcheprova, Z., Baldi, S., Frei, S.B., Gonnet, G., and Barral, Y. (2008). A mechanism for asymmetric segregation of age during yeast budding. Nature 454, 728–734.

61. Sinclair, D.A., Mills, K., and Guarente, L. (1998). Molecular mechanisms of yeast aging. Trends Biochem Sci 23, 131–134.

62. Singh, P., Ramachandran, S.K., Zhu, J., Kim, B.C., Biswas, D., Ha, T., Iglesias, P.A., and Li, R. (2017). Sphingolipids facilitate age asymmetry of membrane proteins in dividing yeast cells. Mol Biol Cell 28, 2712–2722.

63. Smith, S.E., Rubinstein, B., Mendes Pinto, I., Slaughter, B.D., Unruh, J.R., and Li, R. (2013). Independence of symmetry breaking on Bem1-mediated autocatalytic activation of Cdc42. J Cell Biol 202, 1091–1106.

64. Spokoini, R., Moldavski, O., Nahmias, Y., England, J.L., Schuldiner, M., and Kaganovich, D. (2012). Confinement to organelle-associated inclusion structures mediates asymmetric inheritance of aggregated protein in budding yeast. Cell Rep 2, 738–747.

65. Tarassov, K., Messier, V., Landry, C.R., Radinovic, S., Serna Molina, M.M., Shames, I., Malitskaya, Y., Vogel, J., Bussey, H., and Michnick, S.W. (2008). An in vivo map of the yeast protein interactome. Science 320, 1465–1470.

66. Thieleke-Matos, C., Osorio, D.S., Carvalho, A.X., and Morais-de-Sa, E. (2017). Emerging Mechanisms and Roles for Asymmetric Cytokinesis. Int Rev Cell Mol Biol 332, 297–345.

67. Toenjes, K.A., Simpson, D., and Johnson, D.I. (2004). Separate membrane targeting and anchoring domains function in the localization of the S. cerevisiae Cdc24p guanine nucleotide exchange factor. Curr Genet 45, 257–264.

68. Tolliday, N., Pitcher, M., and Li, R. (2003). Direct evidence for a critical role of myosin II in budding yeast cytokinesis and the evolvability of new cytokinetic mechanisms in the absence of myosin II. Mol Biol Cell 14, 798–809.

69. Tu, B.P., Ho-Schleyer, S.C., Travers, K.J., and Weissman, J.S. (2000). Biochemical basis of oxidative protein folding in the endoplasmic reticulum. Science 290, 1571–1574.

70. van Drogen, F., and Peter, M. (2002). Spa2p functions as a scaffold-like protein to recruit the Mpk1p MAP kinase module to sites of polarized growth. Curr Biol 12, 1698–1703.

71. Verna, J., Lodder, A., Lee, K., Vagts, A., and Ballester, R. (1997). A family of genes required for maintenance of cell wall integrity and for the stress response in Saccharomyces cerevisiae. Proceedings of the National Academy of Sciences of the United States of America 94, 13804–13809.

72. Versele, M., and Thorner, J. (2004). Septin collar formation in budding yeast requires GTP binding and direct phosphorylation by the PAK, Cla_4_. Journal of Cell Biology 164, 701–715.

73. Versele, M., and Thorner, J. (2005). Some assembly required: yeast septins provide the instruction manual. Trends Cell Biol 15, 414–424.

74. Voth, W.P., Olsen, A.E., Sbia, M., Freedman, K.H., and Stillman, D.J. (2005). ACE2, CBK1, and BUD4 in budding and cell separation. Eukaryot Cell 4, 1018–1028.

75. Watts, F.Z., Shiels, G., and Orr, E. (1987). The yeast MYO1 gene encoding a myosin-like protein required for cell division. EMBO J 6, 3499–3505.

76. Weirich, C.S., Erzberger, J.P., and Barral, Y. (2008). The septin family of GTPases: architecture and dynamics. Nat Rev Mol Cell Biol 9, 478–489.

77. Weiss, E.L. (2012). Mitotic exit and separation of mother and daughter cells. Genetics 192, 1165–1202.

78. Wloka, C., and Bi, E. (2012). Mechanisms of cytokinesis in budding yeast. Cytoskeleton (Hoboken) 69, 710–726.

79. Wu, C.F., Chiou, J.G., Minakova, M., Woods, B., Tsygankov, D., Zyla, T.R., Savage, N.S., Elston, T.C., and Lew, D.J. (2015). Role of competition between polarity sites in establishing a unique front. Elife 4.

80. Wu, N., Sarna, L.K., Hwang, S.Y., Zhu, Q., Wang, P., Siow, Y.L., and O, K. (2013). Activation of 3-hydroxy-3-methylglutaryl coenzyme A (HMG-CoA) reductase during high fat diet feeding. Biochim Biophys Acta 1832, 1560–1568.

